# Monocarboxylate transporter 1 in Schwann cells is critical for maintenance of sensory nerve myelination during aging

**DOI:** 10.1101/686832

**Authors:** Mithilesh Kumar Jha, Youngjin Lee, Katelyn A. Russell, Fang Yang, Raha M. Dastgheyb, Pragney Deme, Xanthe Heifetz Ament, Weiran Chen, Ying Liu, Yun Guan, Michael J. Polydefkis, Ahmet Hoke, Norman J. Haughey, Jeffrey D. Rothstein, Brett M. Morrison

**Affiliations:** Department of Neurology, Johns Hopkins University School of Medicine, Baltimore, MD 21205, United States; Department of Biomedical Sciences, City University of Hong Kong, Hong Kong SAR, Hong Kong; Department of Anesthesiology and Critical Care Medicine, Johns Hopkins University School of Medicine, Baltimore, MD 21205, United States; Department of Neurological Surgery, Johns Hopkins University School of Medicine, Baltimore, MD 21205, United States

**Keywords:** lactate, monocarboxylate transporter, MCT1, Schwann cell, myelination, sensory axons, peripheral nerve, metabolism, triacylglycerides

## Abstract

Schwann cell (SC)-specific monocarboxylate transporter 1 (MCT1) knockout mice were generated by mating MCT1*^f/f^* mice with myelin protein zero (P0)-Cre mice. P0-Cre^+/-^, MCT1*^f/f^* mice have no detectable early developmental defects, but develop hypomyelination and reduced conduction velocity in sensory, but not motor, peripheral nerves during maturation and aging. Furthermore, enlarged node length and reduced mechanical sensitivity were evident in aged P0-Cre^+/-^, MCT1*^f/f^* mice. MCT1 deletion in SCs impairs both their glycolytic and mitochondrial functions, leading to altered lipid metabolism of triacylglycerides, diacylglycerides, and sphingomyelin, decreased expression of myelin-associated glycoprotein (MAG), and increased expression of c-Jun and p75-neurotrophin receptor, suggesting a regression of SCs to a less mature developmental state. Taken together, our results define the essential role of SC MCT1 in both SC metabolism and peripheral nerve maturation and aging.

**Main Points:**

- SC MCT1 deficiency causes hypomyelination of sensory, but not motor, axons during aging
- Enlarged node length of sensory axons is evident in mutant mouse with SC-specific MCT1 deletion
- Selective ablation of MCT1 within SCs impairs glycolytic and mitochondrial functions
- SC-specific MCT1 deficiency impairs proteins that regulate myelin and lipid metabolism in peripheral nerves

## 1. INTRODUCTION

Structural maintenance and proper functioning of the peripheral nervous system (PNS) require high metabolic supply. Aging peripheral nerves have progressive reduction in energy stores (Low, Schmelzer, & Ward, 1986) as well as blood flow due to reduction of microvasculature caliber (Kihara, Nickander, & Low, 1991). Although Schwann cells (SCs) play a critical role in myelinating axons of peripheral nerves and facilitating saltatory conduction, growing evidence suggests that similar to oligodendrocytes and astrocytes in the central nervous system (CNS) (Funfschilling et al., 2012; Lee et al., 2012; Pellerin & Magistretti, 1994), SCs may also be a primary metabolic supporter to axons in the PNS (Beirowski et al., 2014; Vega, Martiel, Drouhault, Burckhart, & Coles, 2003). Such metabolic support is quite important for maintenance of axonal integrity and function in the PNS, where there are long axons that have distal regions relatively isolated from their cell bodies. Emerging evidence outlines the critical role of the axon-SC relationship and associated bioenergetic crosstalk in peripheral nerve health and diseases. Transfer of metabolic substrates from SCs to axons has been previously reported in rodent models (Vega et al., 2003), and SC metabolism has been found to be critical for developmental myelination, myelin maintenance, and support of axon integrity and function (Beirowski et al., 2014; Viader et al., 2011; Yin et al., 2016).

Uninterrupted energy supply is necessary for the survival of peripheral axons. The presence of glycogen, which is the primary intracellular metabolic storage molecule, within SCs establishes this cell type as an important source of energy substrates, primarily in the form of lactate, for myelinated axons during aglycemia (Brown, Evans, Black, & Ransom, 2012). Growing evidence suggests a critical role of lactate as a dynamic metabolite in health and diseases of peripheral nerves (Beirowski et al., 2014; Jha & Morrison, 2018). During catabolic processes, glycogen present in SCs is metabolized into lactate that may be transported to axons via monocarboxylate transporters (MCTs), which are bi-directional transporters of protons and monocarboxylates, including lactate, across plasma membranes. Within axons, lactate or other monocarboxylates may then be oxidized to pyruvate and metabolized through the Krebs cycle for ATP production (Beirowski et al., 2014; Brown et al., 2012). MCT1 is the most abundantly expressed lactate transporter in peripheral nerves, being expressed in SCs along with other cell types (Domenech-Estevez et al., 2015; Morrison et al., 2015; Takebe, Nio-Kobayashi, Takahashi-Iwanaga, & Iwanaga, 2008). We have previously reported that MCT1 is critical for nerve regeneration following injury, and this altered regeneration is potentially due to impairment of intrinsic SC biology necessary for proliferation, de-differentiation, or cytokine release (Morrison et al., 2015). Alternatively, reduced nerve regeneration may be due to altered metabolic support from SCs to regenerating axons. Though not yet specifically investigated in SCs *in vivo*, oligodendrocyte progenitor cells (OPCs) in the CNS utilize lactate through MCTs to promote proliferation and differentiation (Ichihara et al., 2017). Surprisingly, downregulation of MCT1 in SCs *in vitro* by lentiviral shRNAs reportedly increases myelination in DRG neuron/SCs myelinating co-cultures (Domenech-Estevez et al., 2015). Hence, the exact role of MCT1 expressed in SCs in peripheral nerve biology is not yet clear.

To directly interrogate the role of MCT1 in SCs *in vivo*, we have developed P0-Cre^+/-^, MCT1*^f/f^* mice (P0-cMCT1 KO), which have SC-specific ablation of MCT1, by mating newly generated MCT1*^f/f^* mice with myelin protein zero Cre-recombinase (P0-Cre) mice. In this study, we evaluate the integrity of peripheral nerve axons and myelin, function of peripheral nerves by both electrophysiologic measurements and behavior, and characterize the specific metabolic alterations in P0-cMCT1 KO mice during development and aging. We find that SC-specific MCT1 deletion impairs glycolytic and mitochondrial functions, depletes the nerve of critical lipids, and results in hypomyelination and functional deficits in sensory, but not motor, peripheral nerves during aging.

## 2. MATERIALS AND METHODS

### 2.1 Generation and Propagation of Genetically Modified Mice

All animal experiments were carried out in compliance with the protocols approved by the Johns Hopkins University Institutional Animal Care and Use Committee (IACUC). SC-specific MCT1 knockout mice (P0-Cre^+/-^, MCT1*^f/f^* or noted as P0-cMCT1 KO mice) and control littermates (P0-Cre^-/-^, MCT1*^f/f^* or noted as wild-type [WT]) were generated by crossing MCT1*^f/f^* mice to transgenic mice with Cre recombinase driven by myelin protein zero (P0) promoter (P0-Cre mice, stock no. 017927, The Jackson Laboratory) (Feltri et al., 1999). To generate the MCT1*^f/f^*, an original targeting construct for MCT1 exon2, containing ATG translation initiation site, was obtained from the European Conditional Mouse Mutagenesis Program (EUCOMM) (Skarnes et al., 2011). 5’ arm of the targeting construct was modified for improving the efficiency of homologous recombination. Germline transmission for the chimeric mice, produced by targeting 129 ES cells with the construct, was confirmed by a series of genotyping PCR analyses and several transgenic lines were established. To remove the selection cassette from the targeting construct, the transgenic mice were bred with FLP recombinase deleter mice (Stock 009086; Jackson Laboratory, Bar Harbor ME). The resulting MCT1*^f/f^* mice, harboring conditional/inducible-ready alleles, were back-crossed for two generations with wild-type C57/BL6 mice. For all comparisons, only littermate controls were used. Validation for MCT1*^f/f^* mice was conducted by Southern blot and PCR. Genotypes for MCT1*^f/f^* and P0-cMCT1 KO mice were performed by PCR with the following primer pairs: MCT1*^f/f^* forward, 5’-GCA GCA TGT GGT CCT CTC TTA AG-3’, reverse 5’-GTC CTC ACC TCT CTG TGC-3’; MCT1, forward, 5’-GCA GCA TGT GGT CCT CTC TTA AG-3’, reverse, 5’-TGG TTC TCT TGT TAT CAG TGT TGG GTG-3’; P0-Cre, forward, 5’-GTG AAA CAG CAT TGC TGT CAC TT-3’, reverse, 5’-GCG GTC TGG CAG TAA AAA CTA TC-3’. PCR amplification (using Radiant™ 2x RED Taq Mix; Alkali Scientific) was performed in 39 cycles (for MCT1*^f/f^* and MCT1) or 35 cycles (for P0-Cre), with each cycle consisting of denaturation at 94°C for 30 sec, annealing at 52 to 65°C for 30-45 sec, depending on the primer pair’s annealing temperature, and extension at 72°C for 45-60 sec.

### 2.2 Nerve Histology and Morphometry

Deeply anesthetized mice were transcardially perfused with 0.1 M phosphate buffer saline (PBS) followed by 4% paraformaldehyde fixative and nerves or dorsal/ventral roots of spinal nerves were dissected. Tissue for immunofluorescence staining was post-fixed for 4 h in 4% paraformaldehyde, cryoprotected in 25% sucrose and sectioned on a Leica CM1900 cryostat. Sections (20 μm thickness) were immunostained on slides for MCT1 (generated for laboratory (Morrison et al., 2015); 1:200), S100 (Dako; 1:500), phosphorylated neurofilament (SMI31; Covance; 1:5000), c-Jun (clone 60A8; Cell Signaling Technology; 1:300) or CD271 (p75^NTR^; BioLegend, 1:2000) either alone or in combination. Photomicrographs were acquired with ZEN Digital Imaging for LSM 800 (Zeiss). Tissue for electron microscopy or semi-thin nerve histology was post-fixed with 2.5% glutaraldehyde in 4% paraformaldehyde for at least 3 days and embedded in Epon 812 resin. Embedded nerves were cut either semi-thin (1 μm) and stained with toluidine blue or thin (70 nm) and stained with citrate/uranyl acetate. Toluidine blue stained sections were used for quantification of myelinated axon number, myelinated axon diameter, myelin thickness, or *g* ratio. Our technique is well supported in the literature, as an identical procedure has been followed in a number of publications (Painter et al., 2014; Viader et al., 2013). For each of these histologic features, photomicrographs of nerves or roots were taken on Nikon E800, imported and quantified manually with Zeiss AxioVision 4.8 software. *g* ratio was calculated as the ratio of the diameter of axons divided by the diameter of myelin sheaths. If more than one fascicle was present in a sample, the largest nerve fascicle was quantified in its entirety. An experimenter, who was blinded to animal genotypes, performed all morphometric analyses.

### 2.3 RNA Preparation and Quantitative Real-Time Reverse Transcription-PCR

Deeply anesthetized mice were transcardially perfused with 0.1 M PBS to remove the blood, and the sciatic nerves or dorsal and ventral roots of spinal nerves were rapidly dissected. RNA was isolated by an RNeasy Mini Kit (Qiagen), reverse transcribed to cDNA with a High Capacity cDNA Reverse Transcription Kit (Applied Biosystems) and quantified by real-time RT PCR using Taqman probes (Applied Biosystems) for MCT1 (Thermo Fisher Scientific; Catalog # 4351372), MCT2 (Thermo Fisher Scientific; Catalog # 4331182), MCT4 (Thermo Fisher Scientific; Catalog # 4331182), GLUT1 (Thermo Fisher Scientific; Catalog # 4331182), GLUT3 (Thermo Fisher Scientific; Catalog # 4331182), TRPV1 (Thermo Fisher Scientific; Catalog # 4448892), or GAPDH (Thermo Fisher Scientific; Catalog # 4352339E) on a StepOne Plus RT-PCR System (Applied Biosystems).

### 2.4. Western Blotting

Peripheral nerves dissected after transcardial perfusion of deeply anesthetized mice with

0.1 M PBS were homogenized in T-PER (Thermo Scientific) and run on Mini-Protean TGX Gels (10%; Bio-Rad) and transferred to nitrocellulose membranes (Bio-Rad). For all Western blots, 15-30 μg of proteins were separated on the gel. Membranes were incubated overnight with MCT1 (generated for laboratory (Morrison et al., 2015); 1:200), P0 (Aves Labs. Inc.; catalog # PZO; myelin protein-zero chicken polyclonal anti-peptide antibody; 1:4000), MBP (Millipore Sigma; catalog # AB980; anti-myelin basic protein antibody; 1:250), or MAG (Millipore Sigma; catalog # PA5-30087; polyclonal antibody; 1:500) antibodies and visualized with Amersham ECL Reagant (GE Healthcare) on ImageQuant LAS 4000 (GE Healthcare). After visualizing for above primary antibodies, blots were stripped with Restore Western Blot Stripping Buffer (Thermo Scientific), reprobed overnight with β-actin (Millipore Sigma; catalog # A5316; monoclonal anti-β-actin antibody; 1:5000), and again visualized by ECL reagent, as described above.

### 2.5 Nerve Conduction Studies

Electrophysiological recordings were performed to measure sensory nerve action potentials (SNAPs) from the tail nerve and compound muscle action potentials (CMAPs) by using a Neurosoft-Evidence 3102evo electromyograph system (Schreiber & Tholen Medizintechnik, Stade, Germany). During all recording sessions, mice were anesthetized with 2% isoflurane and positioned face down. To accurately localize testing sites for SNAP recordings, a baseline site between the 2nd and 3rd vertebrae (at top/beginning of the tail) was established and electrodes (30G platinum subdermal needle electrodes, Natus Medical Inc., San Carlos, CA) were placed at 5 mm and 10 mm (+/- recording), 20 mm (ground/reference), 30 mm, 35 mm (+/-proximal stimulation), 45 mm, and 50 mm (+/-distal stimulation) from the baseline site. Stimulation of each nerve segment was performed, with increasing voltage, until the maximal response was achieved, as evidenced by no further increase or reduction in SNAP amplitude, despite an increase in stimulation voltage. Response latency for each proximal or distal stimulation was measured from stimulus onset, and peak-to-peak amplitudes were calculated. Sensory NCV was calculated by dividing the distance between the points of proximal and distal stimulations (here, fixed at 15 mm) by the difference between their response latencies. The temperature of the tail was measured during each experiment by the Digi-Sense Infrared Thermometer (Model 20250-05). The CMAPs were determined by placing stimulating electrodes (27G stainless steel needle electrodes, Natus Medical Inc., San Carlos, CA) at the sciatic notch and Achilles tendon, and recording electrodes in the lateral plantar muscles of the foot. Similar to the SNAP recordings, stimulation voltage for CMAP recordings were adjusted for maximal response. Nerves were stimulated by very short (< 0.2 millisecond) electrical impulses. Response latency for each proximal or distal stimulation was measured from stimulus onset, and peak-to-peak amplitudes were calculated. Motor NCV was calculated by dividing the distance between sciatic notch and Achilles tendon by the difference between the response latencies. Unless otherwise indicated, all nerve conduction studies were conducted at room temperature. The investigator performing electrophysiological recordings was blinded to mice genotypes throughout the study.

### 2.6 Measurement of Node of Ranvier length

The sural nerve from aged mice was dissected immediately, desheathed, and dissociated into single fibers by treatment with collagenase (3.6 mg/ml; Sigma) for 1 h at room temperature (RT). Fibers were teased apart gently and spread on coverslips that had been coated in a few spots with Cell-Tak (Collaborative Research, Bedford, MA). After teasing, the preparations were fixed in 4% paraformaldehyde in 0.1 M phosphate buffer, pH 7.2, for 30 min, rinsed in 0.05 M phosphate buffer, pH 7.4, for 10 min, and air-dried. Teased nerve fibers were immunostained for anti-Caspr antibody (rabbit polyclonal; 1:200; Abcam) and anti-Ankyrin-G antibody (mouse monoclonal; 1:200; clone N106/36; NeuroMab). Images were acquired with ZEN Digital Imaging for LSM 800 (Zeiss) and analyzed by using ZEN blue edition software (Zeiss). The node length was measured without correcting for fiber shrinkage during fixation.

### 2.7 Intra-epidermal Nerve Fiber Analysis

Analysis of footpads for intra-epidermal nerve fiber density (IENFD) was performed as previously described (Chiorazzi et al., 2018). Briefly, footpads were dissected out and were washed in 0.1 M phosphate buffer and placed in cryoprotectant solution (30% glycerol). Tissue blocks were sectioned by freezing microtome at 50 μm intervals and immunohistochemical staining was performed for protein gene product 9.5 (PGP 9.5; rabbit polyclonal; AbD Serotec, a Bio-Rad Company, Kidlington, UK) using a free-floating protocol. The total number of PGP 9.5-positive IENF in each section was counted, the length of the epidermis was measured and the linear density of IENF/mm was obtained. An experimenter, who was blinded to animal genotype, performed the tissue processing, immunostaining, and analysis.

### 2.8 Behavioral Studies

An experimenter blinded to animal genotype carried out all behavioral tests. One day before behavioral testing, mice were acclimated to the environment for 1 h. Mechanical sensitivity was assessed with the von Frey test by the frequency method. Two calibrated von Frey monofilaments (low force, 0.07 g; high force, 0.45 g) were used. Each von Frey filament was applied perpendicularly to the plantar side of hind paw for ∼1 s; the stimulation was repeated 10 times to the paw. The occurrence of paw withdrawal in each of these 10 trials was expressed as a percent response frequency: paw withdrawal frequency = (number of paw withdrawals/10 trials) × 100%. Mean of the left and the right hindpaw measurements from each mouse were calculated. For the Hargreaves test, mice were placed under a transparent plastic box (4.5 cm × 5 cm × 10 cm) on a glass floor. Infrared light was delivered through the glass floor to the hind paw. After acclimatization sessions, the latency for the animal to withdraw its hind paw was measured.

### 2.9 SC Culture and Seahorse Bioenergetic Analysis

Cultures of SCs were prepared from the sciatic nerves of 4-week-old mice as described previously (Scheib & Hoke, 2016) with slight modifications. Briefly, dissected sciatic nerves were cut into small pieces and dissociated at 37°C in 0.3% collagenase (Gibco) for 45 minutes and 0.25% trypsin for 15 minutes. Dissociated cells were resuspended in SC media (ScienceCell), which consists of basal media supplemented with 5% FBS, 1% SC growth supplement and 1% penicillin/streptomycin solution, and plated. On day-one after plating, the media was switched to SC purification media, which consists of SC media supplemented with 10 µM arabinosylcytosine (AraC, Millipore Sigma), for two days. The media was then switched to SC media for two days, which was followed by switching to SC purification media for the next two days. SCs purity (greater than 95%) was confirmed by immunostaining with antibody against S100 and were used for measurements of oxygen consumption and extracellular acidification using XF96 extracellular flux analyzer (Seahorse Bioscience, North Billerica, MA) (Yang et al., 2013) following the manufacturer’s instructions. SCs were grown in the Seahorse XF96 cell culture microplate at 10,000 cells/well. Bioenergetic analysis was performed by sequentially injecting 2 μM oligomycin, 4 μM carbonyl cyanide *p*-(trifluoromethoxy) phenylhydrazone (FCCP), 0.5 μM rotenone and 4 μM antimycin. Data are expressed as oxygen consumption rate (OCR) in picomole per minute and extracellular acidification rate (ECAR) in milli pH per minute for 10,000 cells.

### 2.10 Nerve Lipidomics

Sural nerves from aged (eight months old) mice were dissected, cleaned from surrounding fat and connective tissue, and snap frozen using dry ice. Nerve tissues were extracted using a modified Bligh and Dyer procedure to obtain a crude lipid fraction (Bligh & Dyer, 1959). In brief, tissues were homogenized in ddH_2_0 and protein normalized volumes of tissue homogenate were gently mixed in a glass vial with 940ml ddH_2_O and 2.9mL methanol/dichloromethane (2:0.9, v/v) containing the following internal standards: N-lauroyl-D-erythro-sphingosine (Cer d18:1/12:0, 6 ng/mL), 1,3(d5)-dihexadecanoyl-glycerol (d5-DAG d16:0/16:0, 12.5 ng/mL), D-galactosyl-β-1,1’ N-lauroyl-D-erythro-sphingosine (GlcCer d18:1/12:0, 3.3 ng/mL), D-lactosyl-β-1,1’ N-lauroyl-D-erythro-sphingosine (LacCer 18:1/12:0, 10.6 ng/mL), 1,3(d5)-dihexadecanoyl-2-octadecanoyl-glycerol (D-5 TAG 16:0/18:0/16:0, 0.5 ng/mL), cholesteryl-d7 palmitate (cholesteryl-d7 ester 16:0, 30 ng/mL), 1,2-dilauroyl-sn-glycero-3-phosphate (sodium salt) (PA d12:0/12:0, 1025 ng/mL), 1,2-dilauroyl-sn-glycero-3-phosphocholine (PC 12:0/12:0, 0.2 ng/mL), 1,2-dilauroyl-sn-glycero-3-phosphoethanolamine (PE d12:0/12:0, 1.6 ng/mL), 1,2-dilauroyl-sn-glycero-3-phospho-[1’-rac-glycerol] (PG d12:0/12:0, 200 ng/mL), 1,2-dilauroyl-sn-glycero-3-phospho-L-serine (PS d12:0/12:0), and N-lauroyl-D-erythro-sphingosylphosphorylcholine (SM d18:1/12:0, 0.3 ng/mL). All internal standards were purchased from Avanti Polar Lipids, Inc. (Alabaster, AL). To obtain a biphasic mixture, an additional 1mL of ddH_2_O and 0.9mL dichloromethane was added and vortexed. The resultant mixture was incubated on ice for 30 min and centrifuged (10 min, 3000g, 4°C) to separate the organic and aqueous phases. The organic phase was removed and stored at −20°C. Just prior to analysis, 1 mL of the organic layer was dried using a nitrogen evaporator (Organomation Associates, Inc., Berlin, MA, USA) and re-suspended in 250µl of running solvent (dichloromethane:methanol (1:1) containing 5mM ammonium acetate), and 5mg/mL of ceramide C17:0 used to track instrument performance. All solvents used were HPLC grade. Samples were analyzed on a 5600 Triple TOF^TM^ mass spectrometer (AB Sciex, Concord, Canada). For data analysis, lipid peak intensities were first log transformed to achieve a normal distribution. They were then filtered based on the following criteria: skewness <10, kurtosis <10, and IQR:standard deviation ratio differing less than 50% from 1.35 (Rice, 2007). Using these criteria 38 of the 755 lipid analytes were excluded from further analyses. Log-transformed lipid intensities were investigated in a univariate analysis, and the means between the two groups were compared using t-tests with a Benjamini-Hochberg correction (Benjamini & Hochberg, 1995) for multiple comparisons (MathWorks, 2018). For visualization, lipid species were Z-scored to have a mean of 0 and a standard deviation of 1. Principal component analysis (PCA) was used to interrogate variations in the lipidomic data. This procedure creates linear combinations of the original variables to create principal components (Jolliffe, 2011) in an unsupervised manner. An overall PCA, which was limited to three components, was conducted on the entire data set. Analysis was conducted using R version 3.5.1 with mixOmics version 3.8 (Rohart, Gautier, Singh, & Le Cao, 2017).

### 2.11 Quantification and Statistical Analysis

Analyses were performed blinded to animal genotype and treatment. Although we did not perform statistical tests to predetermine sample size, our samples sizes are similar to previously published studies in the field. Statistical analyses were performed with GraphPad Prism 8 (GraphPad Software) by using unpaired *t* test with two tails with unequal variance, one-way ANOVA, or two-way ANOVA with post-hoc test when required conditions were met. The number of animals per group or independent repeats (n), the statistical test used for comparison, and the statistical significance (*p* value) was stated for each figure panel in the respective legend. All data were presented as the mean ± SEM unless otherwise noted. Differences in the *p* values of <0.05 were considered statistically significant.

### 2.12 Data Sharing

The data that support the findings of this study are available from the corresponding author upon reasonable request.

## 3. Results

### 3.1 Deletion of MCT1 from SCs reduces its expression in sciatic nerves without any compensatory alterations in expression of other metabolic mediators

With the goal of exploring the role of SC-specific MCT1 in peripheral nerve biology during development and aging, we produced mice with SC-selective ablation of MCT1 (P0-cMCT1 KO) by mating MCT1*^f/f^* mice with P0-Cre mice (**Figure 1a**). P0-Cre mice express Cre-recombinase in both myelinating and non-myelinating SCs without expression in other cells of the peripheral nerve or in oligodendroglia (Feltri et al., 1999). Since MCT1 is expressed in other cells of the peripheral nerve and oligodendroglia (Lee et al., 2012), cell-selective knockout is necessary for defining the function of SC MCT1 on peripheral nerve biology. Photomicrographs of this P0-Cre line mated to YFP reporter mice demonstrate selective expression in SCs (**Figure 1b**), as published previously (Feltri et al., 1999). We also confirmed that the P0-Cre is not itself detrimental for nerve function by measuring sensory and motor nerve conduction velocities (NCV) and tail sensory (sensory nerve action potential [SNAP]), and sciatic nerve motor (compound muscle action potential [CMAP]) amplitudes from P0-Cre^+/-^, MCT1*^wt/wt^* mice (**Supporting Information Figure S1a-d**). We evaluated mice of each genotype for knockout of MCT1 at the levels of protein, by immunofluorescence (**Figure 1c**) and Western blot (**Figure 1d and Supporting Information Figure S7**), and mRNA by real-time RT PCR (**Figure 1e**). Immunofluorescence clearly shows loss of MCT1 from SCs in the endoneurium in P0-cMCT1 KO mice (**Figure 1c; arrows**) without loss of MCT1 from perineurial cells (**Figure 1c; arrowheads**). Ablation of MCT1 in SCs was enough to reduce its expression significantly in whole sciatic nerve (**Figure 1d-e**), suggesting that SCs express relatively high amounts of MCT1 compared to other cells in the peripheral nerve. Interestingly, SC-specific MCT1 deletion did not cause any compensatory alterations in the expression of other metabolic transporters, MCT2, MCT4, GLUT1, and GLUT3, in sciatic nerves (**Figure 1f-i**).

**Figure 1.**
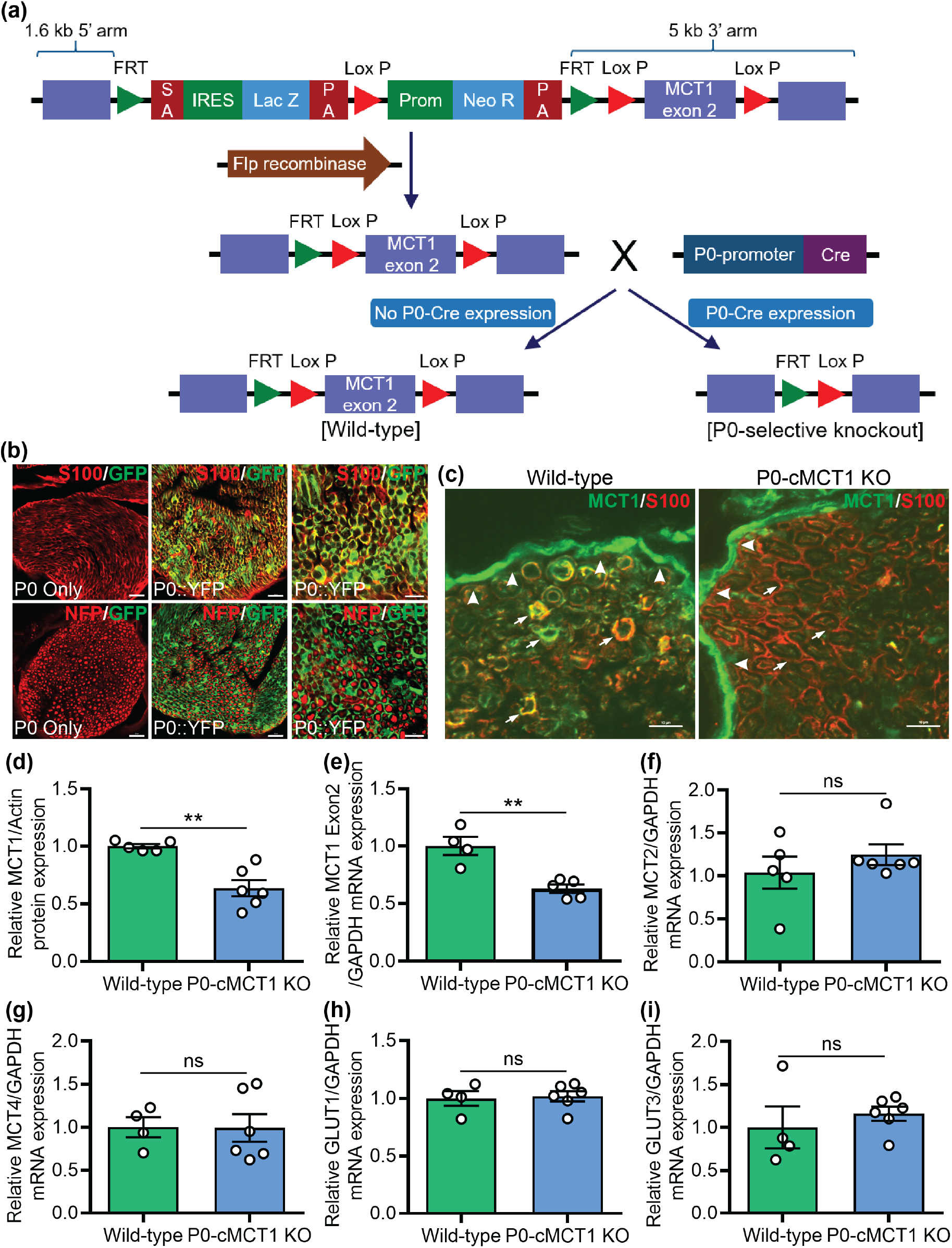
Production and validation of P0-cMCT1 KO mice. (**a**) P0-cMCT1 KO (P0Cre:MCT1*^fl/fl^*) mice were generated by crossing our laboratory-developed MCT1*^f/f^* mice having MCT1 exon2 inserted in between two LoxP sites with P0-Cre transgenic mice. (**b**) P0-Cre^+/-^, RosaYFP mice show complete co-localization of P0-Cre expression (GFP) with S100 (red), a marker for SCs. There is no co-localization of P0-Cre expression (GFP) with phosphorylated neurofilament (NFP; red), which labels axons. Scale bars, 20 µm for left and mid panels; 10 µm for right panel. (**c**) MCT1 immunoreactivity (IR; green) in perineurial cells (arrowheads) and SCs (arrows), co-localized with S100 (red) in wild type nerves (P0:MCT1*^wt/wt^* or wild-type). MCT1 IR (green) is absent from SCs labeled with S100 (red; arrows), but not perineurial cells (arrowheads), in P0-cMCT1 KO mice nerves. Scale bar, 10 µm. (D-I) SC specific MCT1 deletion downregulates the expression of MCT1 protein (**d**) and mRNA (**e**) without altering the expression of MCT2 (**f**), MCT4 (**g**), GLUT1 (**h**), or GLUT3 (**i**) in peripheral nerves of P0-cMCT1 KO mice. Levels of mRNAs are depicted as fold change compared with wild-type mice normalized to their corresponding GAPDH mRNA levels. Mean ± SEM, *n* = 4-6 per group, \*\**p* < 0.01; ns = not significant, unpaired *t* test.

### 3.2 Sensory, but not motor, nerve conduction velocity decreases in maturing P0-cMCT1 KO mice

We measured sensory and motor NCVs and SNAP and CMAP amplitudes to assess peripheral nerve health and axonal integrity during aging following SC-specific MCT1 deletion. Assessment of NCV is an excellent measure of the myelination state of nerves, and SNAP and CMAP amplitudes are reliable indicators of axonal integrity. Although NCV declines with age in humans (Di Iorio et al., 2006), NCV does not decline in male and female mice prior to 20 months of age (Walsh et al., 2015). Moreover, tail SNAP amplitude remains constant over the life span of female mice (Walsh et al., 2015). We evaluated nerve function, by repeated electrophysiologic testing, in P0-cMCT1 KO and littermate control female mice at room temperature, where the tail temperature was usually recorded to be between 22 and 24°C (**Figure 2**). Tail temperature of 4-month-old mice measured at the beginning of electrophysiologic recording was 23.20 ± 0.12 °C for wild-type and 23.59 ± 0.58 °C for P0-cMCT1 KO (Mean ± SEM, *n* = 5 for wild-type and 8 for P0-cMCT1 KO; *p* = 0.533, unpaired *t* test). At 2 months of age, there were no alterations in sensory and motor NCVs (**Figure 2a,c**) or SNAP and CMAP amplitudes (**Figure 2b,d**) following SC-specific MCT1 deletion, suggesting that SC-specific MCT1 has no critical role during this stage of peripheral nerve development. At 4 months of age, however, sensory (**Figure 2a**), but surprisingly not motor (**Figure 2c**), NCV was significantly decreased in mice lacking MCT1 in SCs. The discrepancy in sensory NCV due to MCT1 ablation in SCs was persistent during maturation (3-6 months) and aging (10-12 months) of mice. The decreased sensory NCV and unchanged SNAP amplitude due to MCT1 deletion in SCs at 4 months of age was also confirmed by electrophysiologic recordings performed while maintaining tail temperature at 32–34°C (**Supporting Information Figure S2a,b**). We have also confirmed that the decrease in sensory NCV during aging in female mice is evident in littermate male mice having SC-specific MCT1 deficiency, suggesting that the electrophysiological deficit observed due to MCT1 deficiency in SCs is independent of gender (**Supporting Information Figure S3a,b**). These findings suggest that MCT1 in SCs is crucial for maintenance of sensory nerve function during maturation and aging.

**Figure 2.**
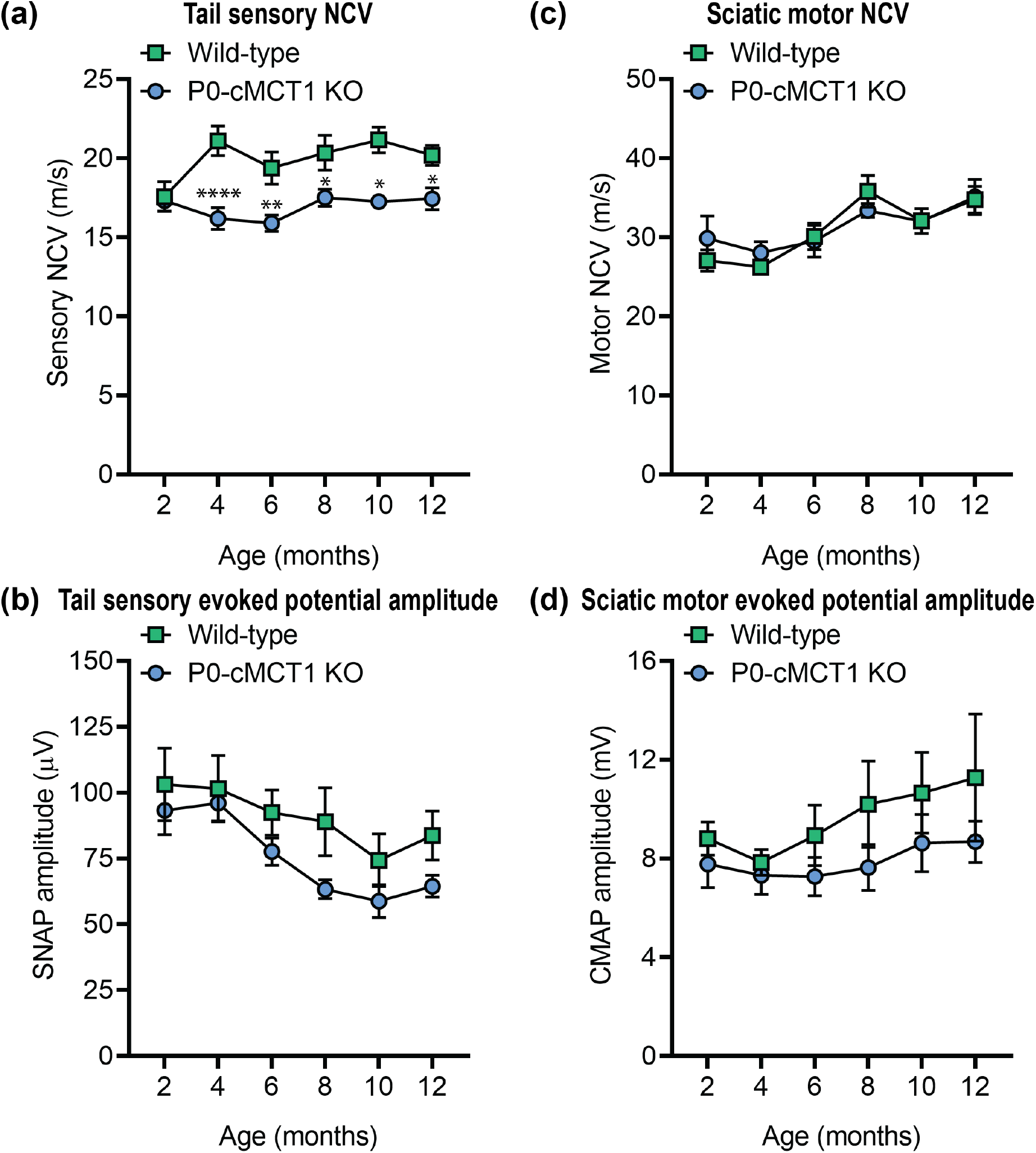
Sensory, but not motor, NCV is decreased in maturing P0-cMCT1 KO mice. (**a** and **c**) SC-specific MCT1 deletion reduces the sensory (**a**), but not motor (**c**), NCV in P0-cMCT1 KO mice, compared to wild-type littermate control mice, by 4 months of age. Mean ± SEM, *n* = 5-7 per group, \**p* < 0.05, \*\**p* < 0.01, \*\*\*\**p* < 0.0001; two-way ANOVA with Holm-Sidak’s multiple comparisons test. (**b** and **d**) No significant impact of SC MCT1 deletion is observed on SNAP (**b**) and CMAP (**d**) amplitudes. The electrophysiological recording was performed at room temperature, where tail temperature was usually recorded to be in between 22 and 24°C. Mean ± SEM, *n* = 5-7 per group. NCV; nerve conduction velocity, SNAP; sensory nerve action potential, CMAP; compound muscle action potential.

### 3.3 SC-specific MCT1 deletion results in sensory nerve hypomyelination during aging

The slower sensory NCV in mice with SC ablation of MCT1 compared with littermate control mice led us to investigate sural nerve (purely sensory) morphology in early postnatal (14-day-old) (**Figure 3a-d** and **Supporting Information Figure S4a-c**), developing (2-month-old) (**Figure 3e-h** and **Supporting Information Figure S4d-f**), maturing (4-month-old) (**Figure 3i-l** and **Supporting Information Figure S4g-i**), and aging (12-month-old) (**Figure 3m-p** and **Supporting Information Figure S4j-l**) mice. Consistent with the electrophysiological recordings of sensory NCV and SNAP amplitudes, we observed a remarkable correlation between genotype and age with respect to myelination and myelinated axon counts. Mice with no MCT1 in SCs developed normally, showed no anatomical difference in term of myelinated axon number, myelination, and myelinated axon caliber at the ages of 14 days and 2 months compared with their littermate control mice (**Figure 3a-h** and **Supporting Information Figure S4a-f**). By 4 months of age, however, the *g* ratio, which reflects the degree of axonal myelination (axon diameter divided by axon plus myelin diameter), for sural nerve axons was significantly increased during maturation, indicating myelin thinning, without any alteration in the number of myelinated axons (**Figure 3k**). The sensory axon hypomyelination in mice having SC-specific MCT1 deficiency was further confirmed by decreased myelin thickness without alterations in axon diameter (**Supporting Information Figure S4g-i)**. Identical to electrophysiology results (**Figure 2**), the hypomyelination by *g* ratio and myelin thickness was present at 12 months, as well (**Figure 3o-p** and **Supporting Information Figure S4j-l**), indicating the crucial role of SC-specific MCT1 in sensory axon myelin maintenance during maturation and aging. For the 4 and 12 month-old sural nerves, the scatter plot of the *g* ratio against the axon diameter revealed that the increased *g* ratio for sensory axons was most prominent for axons with smaller diameters (**Figure 3i,l**). Furthermore, mice with no MCT1 in SCs showed decreased myelinated axon counts during aging that was present by 12 months of age (**Figure 3n**). This was likely due to reduced myelination rather than loss of axons since there was no loss of axons when quantifying all axons greater than 1µm by electron microscopy. The density of axons per 1000 µm^2^ were 21.74 ± 1.73 and 21.98 ± 2.11 for wild-type and P0-cMCT1 KO mice, respectively (11 to 14 random electron micrographs showing a total of 88 to 126 axons per mouse sural nerve captured at x5000 magnification were analyzed for each mouse, Mean ± SEM, n = 3 per group; *p* = 0.935, unpaired *t* test). Though both myelinating and non-myelinating SCs express the P0 promoter (Cheng & Mudge, 1996; Georgiou & Charlton, 1999), we did not see any morphologic abnormalities in Remak bundle organization at any age due to non-myelinating SC MCT1 ablation (**Figure 3**, left panel’s electron micrographs). Besides the overall hypomyelination, alterations in the length of nodes of Ranvier along axons can also contribute to overall NCV. In aged mice, the node length, as measured by distance between paranodes (distance between two Caspr clusters), was increased in sural nerves from P0-cMCT1 KO mice compared to wild-type mice (**Figure 3q,r**). The expansion in nodal length increases nodal capacitance and slows intra-axial current flow from the node into the internodes. These findings demonstrate that MCT1 in SCs is critical for preservation of sensory axon myelin, node length during aging.

**Figure 3.**
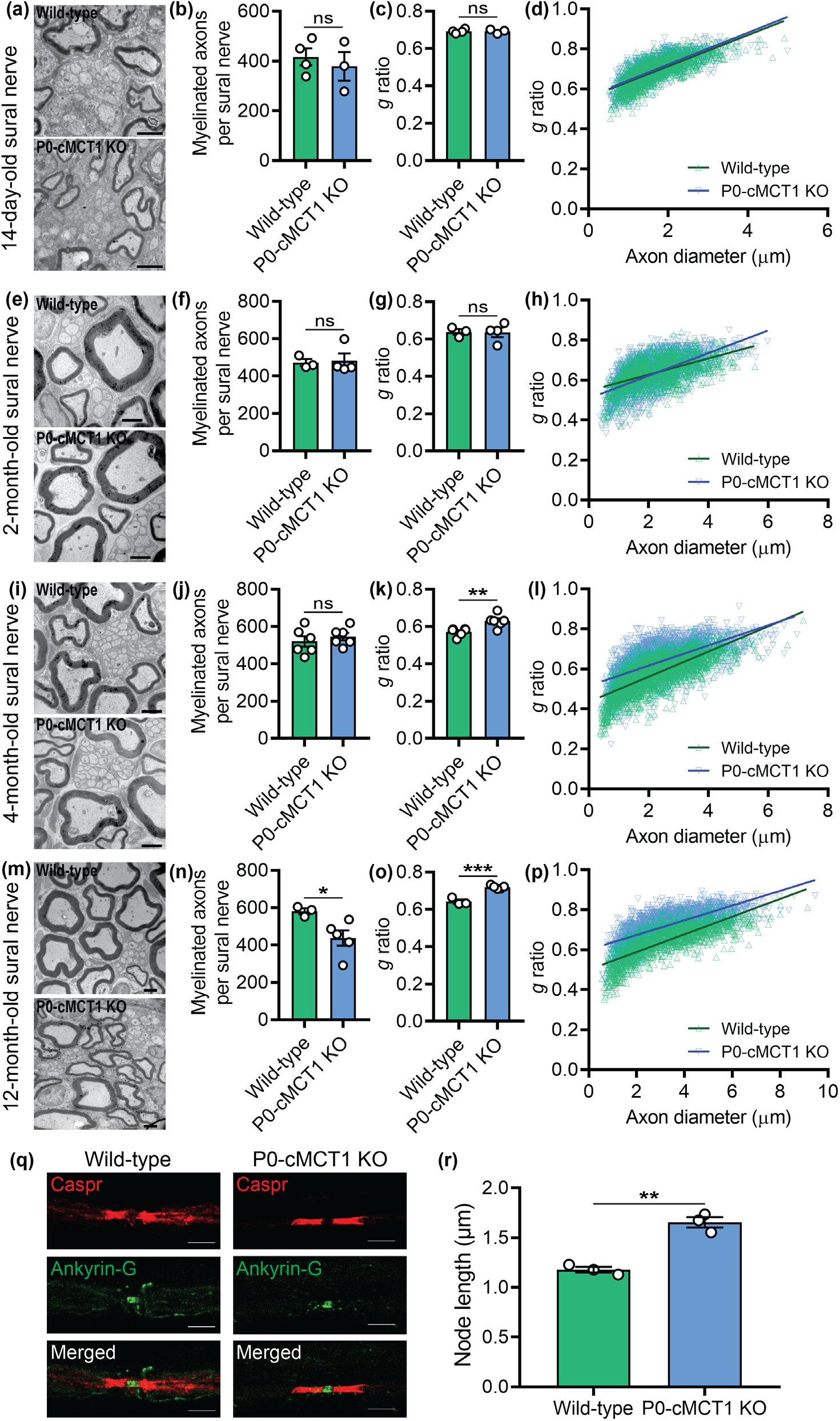
SC-specific MCT1 is essential for maintenance of myelination and nodes of Ranvier. In 14-day-old (**a-d**) and 2-month-old (**e-h**) wild-type and P0-cMCT1 KO mice sural nerves, electron micrographs (**a,e**), myelinated axon count (**b,f**), *g* ratio (**c,g**), and scatter plot graph displaying *g* ratio (y-axis) in relation to axon diameter (x-axis) of individual fiber (**d,h**) confirmed normal embryonic and early post-natal sural nerve development in P0-cMCT1 KO mice. In 4-month-old wild-type and P0-cMCT1 KO mice sural nerves, electron micrographs (**i**), myelinated axon count (**j**), *g* ratio (**k**), and scatter plot graph displaying *g* ratio (y-axis) in relation to axon diameter (x-axis) of individual fiber (**l**) confirmed sural nerve hypomyelination in P0-cMCT1 KO mice. In 12-month-old wild-type and P0-cMCT1 KO mice sural nerves, electron micrographs (**m**), myelinated axon count (**n**), *g* ratio (**o**), and scatter plot graph displaying *g* ratio (y-axis) in relation to axon diameter (x-axis) of individual fiber (**p**) confirmed sural nerve hypomyelination in P0-cMCT1 KO mice. There was also a significant reduction in myelinated axons at 12 months (**n**), but not earlier time points, in P0-cMCT1 KO mice. Mean ± SEM, *n* = 3-6 per group, \**p* < 0.05; \*\**p* < 0.01; \*\*\**p* < 0.001; ns = not significant, unpaired *t* test. Scale bar, 2 µm. The *g* ratio between wild-type (green line) and P0-cMCT1 KO (blue line) was not different in 14-day-old and (**c**) and 2-month-old mice (**d**) but was significantly different in 4-month-old (**h**) and 12-month-old (**l**) mice (for 14-day-old mice: *p* > 0.05; *t* = 0.6309, for 2-month-old mice: *p* > 0.05; *t* = 3.010, for 4-month-old mice: *p* < 0.0001; *t* = 29.29, for 12-month-old mice: *p* < 0.0001; *t* = 30.30, df = 16066, one-way ANOVA with Bonferroni’s multiple comparisons test). (**q** and **r**) Teased sural nerve fibers from 8 month old wild-type and P0-cMCT1 KO mice sural nerves were immunostained using antibodies against Caspr (red; paranodal marker) and Ankyrin G (green; nodal marker) (**q**). Node of Ranvier length was measured as the distance between two paranodal Caspr clusters (**r**). In each mouse, 32-53 nodal gaps were analyzed. Mean ± SEM, *n* = 3 per group, \**p* < 0.05, unpaired *t* test.

The differential sensory and motor electrophysiological features and histological alterations in sural nerve due to SC-specific MCT1 deletion was further confirmed by quantitative assessment of dorsal and ventral roots of the lumbar region of the spinal cord (**Figure 4** and **Supporting Information Figure S5**). Dorsal roots are composed of purely sensory axons, while ventral roots are composed of purely motor axons. Quantitative morphometry of toluidine blue-stained micrographs revealed no alteration in myelinated axon counts and caliber at 12 months of age of both the dorsal and ventral roots due to MCT1 deficiency in SCs (**Figure 4a,b,e,f** and **Supporting Information Figure S5a-f**). However, SC-specific MCT1 deficiency increased the *g* ratio for the dorsal **(Figure 4c,d)**, but not ventral **(Figure 4g,h)**, roots of spinal nerve at 12 months of age, which again appears to primarily affect the smallest myelinated fibers. The reduced myelin thickness of dorsal roots thus confirms the hypomyelination observed more distally in the purely sensory sural nerve.

**Figure 4.**
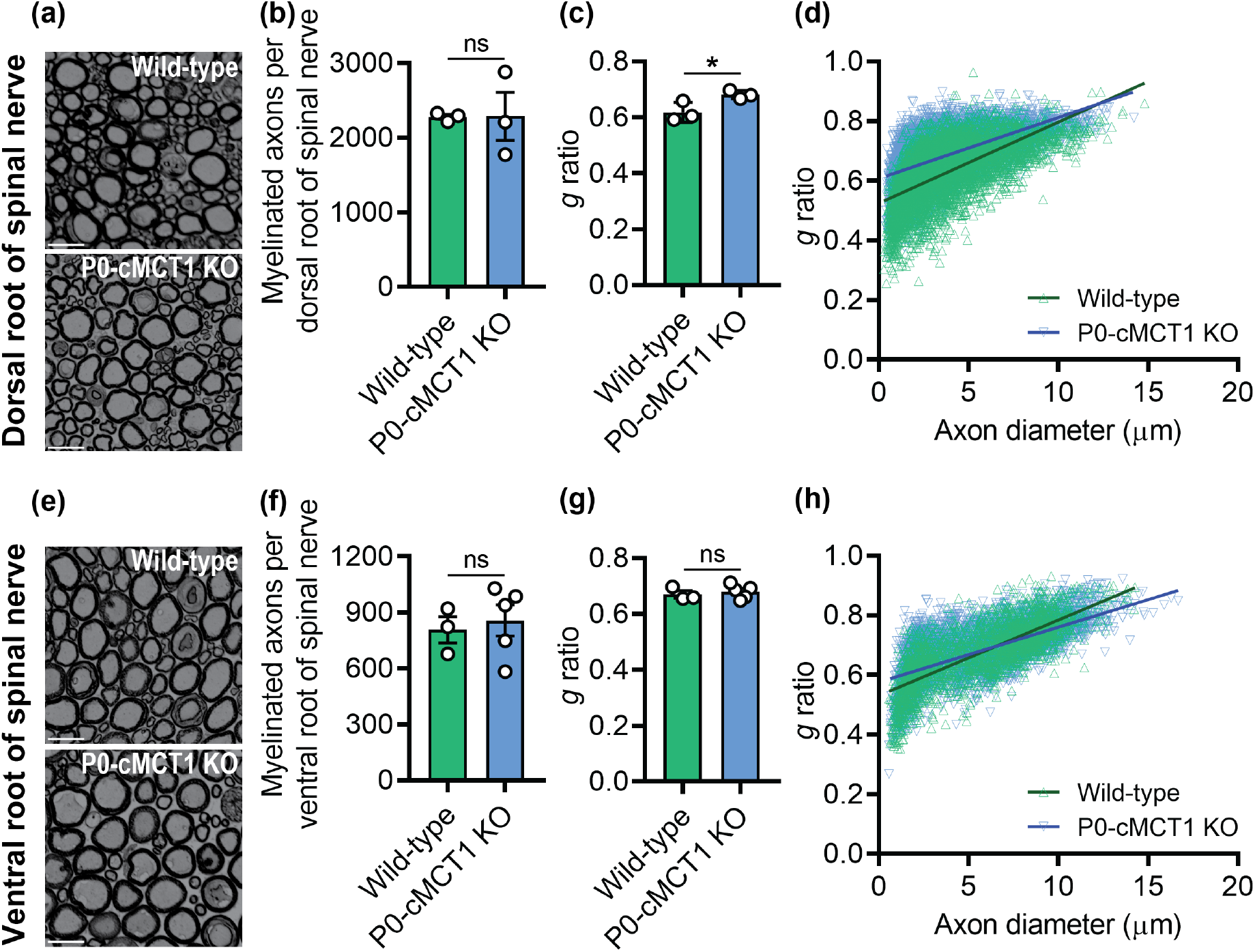
SC-specific MCT1 is critical for myelin maintenance of dorsal, but not ventral, roots of spinal nerves. Photomicrographs (**a**), myelinated axon count (**b**), *g* ratio (**c**), and scatter plot graph displaying *g* ratio (y-axis) in relation to axon diameter (x-axis) of individual fiber (**d**) demonstrated hypomyelination of dorsal spinal nerve roots, without myelinated axon loss, in 12-month old P0-cMCT1 KO mice. Photomicrographs (**e**), myelinated axon count (**f**), *g* ratio (**g**), and scatter plot graph displaying *g* ratio (y-axis) in relation to axon diameter (x-axis) of individual fiber (**h**) confirmed no notable difference between ventral spinal nerve roots of 12-month old P0-cMCT1 KO mice when compared to wild-type mice. Light microscope photomicrographs and subsequent analysis completed on toluidine blue-stained sections. Mean ± SEM, *n* = 3-5 per group, \**p* < 0.05; \*\**p* < 0.01; \*\*\**p* < 0.001; ns = not significant, unpaired *t* test. Scale bar, 20 µm.

**Figure 5.**
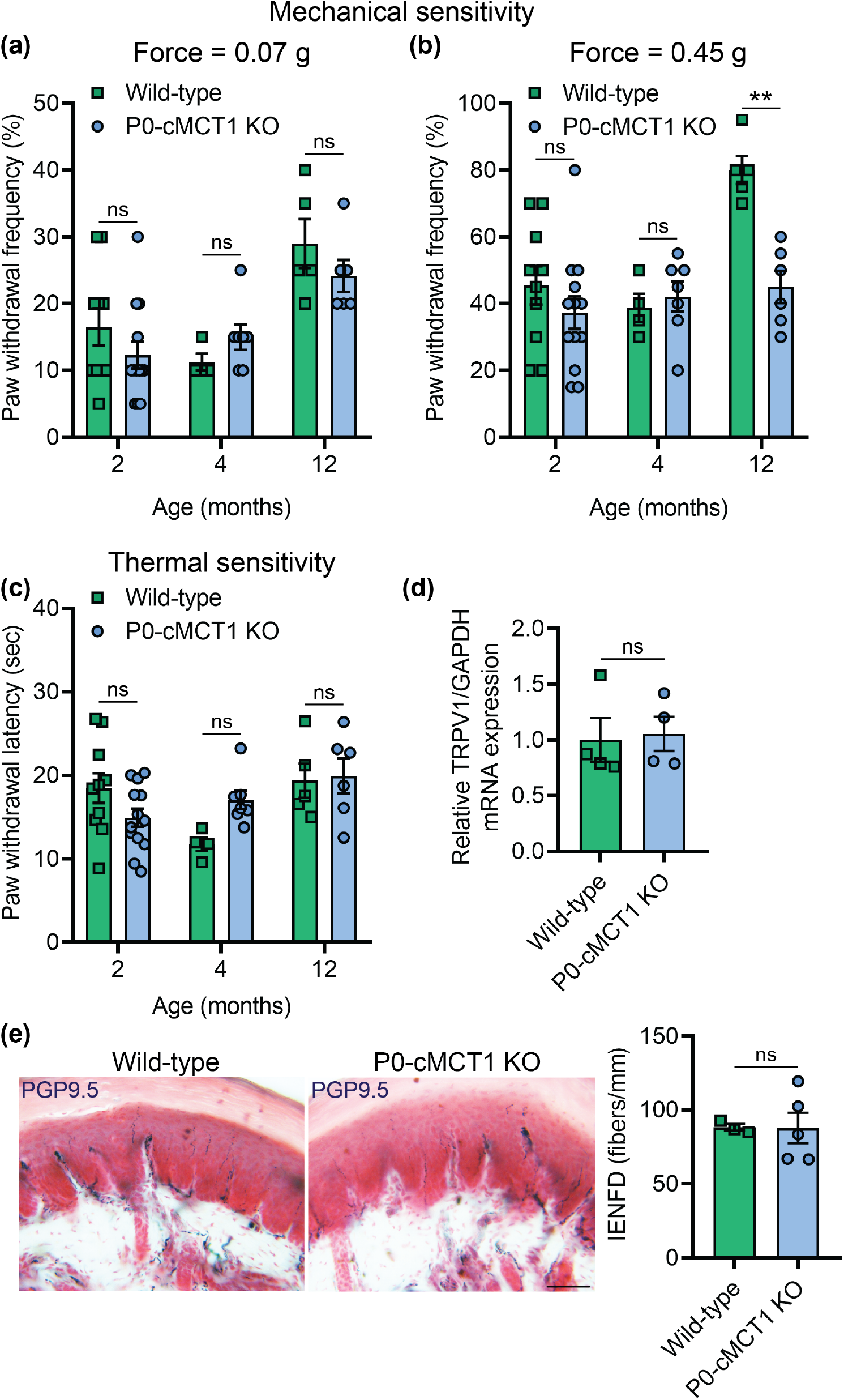
Mechanical sensitivity is impaired in SC-specific MCT1 deficient mice. Assessment of paw withdrawal frequency to mechanical stimulation by calibrated von Frey monofilaments of forces 0.07 g (**a**) and 0.45 g (**b**) in wild-type and P0-cMCT1 KO mice. (**c**) Assessment of thermal sensitivity by using radiant paw-heating assay. Current set at baseline level: 20%, 10-12 sec; cut off time; 30 sec. Mean ± SEM, *n* = 4-10 for wild-type and 6-13 for P0-cMCT1 KO groups, \*\**p* < 0.01; ns = not significant, two-way ANOVA with Bonferroni’s multiple comparisons test. (**d**) Assessment of the TRPV1 mRNA expression by qPCR in sciatic nerves of 4 month old wild-type and P0-cMCT1 KO mice. Levels of TRPV1 mRNA are depicted as fold change compared with wild-type mice normalized to its corresponding GAPDH mRNA levels. *n* = 4 per group, ns = not significant, unpaired *t* test, mean ± SEM. (**e**) Images (left panel) showing representative picture of skin biopsies and quantification of IENFD (right panel) obtained from the footpads of 4-month-old wild-type and P0-cMCT1 KO mice following immunohistochemical staining for PGP9.5. *n* = 3 for wild-type and 5 for P0-cMCT1 KO groups, ns = not significant, unpaired *t* test, mean ± SEM. Scale bar, 50 µm.

### 3.4 Vulnerability of sensory axons is not due to differential expression of MCT1 in sensory nerves compared to motor nerves

The selective impact of reducing SC MCT1 on sensory nerves, with sparing of motor nerves, led us to speculate that there may be differential expression of MCT1 in SCs associated with sensory and motor nerves. To address this issue, we measured the expression of MCT1 mRNA in purely sensory dorsal roots and purely motor ventral roots of spinal nerves. Interestingly, no difference in MCT1 mRNA expression was detected between dorsal and ventral roots (**Supporting Information Figure S5g**), suggesting that the vulnerability of sensory, but not motor, nerves is not due to differential expression of MCT1.

### 3.5 Mice with SC MCT1 deletion develop decreased mechanical sensitivity during aging

Impaired sensory nerve electrophysiology and histology in mice with SC-specific MCT1 deletion led us to investigate if there were sensory deficits in these mice. The behavioral responses to repetitive punctate mechanical stimulation and noxious thermal stimulation were assessed with the von Frey test by the frequency method and the Hargreaves test, respectively **(Figure 5)**. Mice with or without SC MCT1 deletion revealed no difference in mechanical and thermal sensitives during development and maturation (2 and 4 months of age, respectively; **Figure 5a-c**). However, mice with SC MCT1 ablation had decreased mechanical sensitivity during aging compared with littermate control mice, as indicated by decreased paw withdrawal frequency to repetitive high force (0.45g) von Frey filament stimulation (**Figure 5b**), at 12 months of age. The increased paw withdrawal frequency to repetitive high force mechanical stimulation in 12 month wild-type mice is consistent with a recent report suggesting an elevated sensitivity to mechanical stimuli with aging (Weyer et al., 2016). In contrast, no alteration was observed in paw withdrawal latency to noxious thermal stimulation at any age (**Figure 5c**). The decrease in mechanical, and not thermal, sensitivity is interesting and expected as axons conveying thermal signals are unmyelinated. Furthermore, the expression of transient receptor potential vanilloid type 1 (TRPV1), which behaves as a molecular integrator of chemical and thermal noxious stimuli in the PNS determining nociceptive responses, was unchanged in the sciatic nerve following SC-specific deletion of MCT1 (**Figure 5d**). The unaltered thermal sensitivity in aging mice was further supported by unchanged intraepidermal nerve fiber density (IENFD), as measured by PGP9.5-immunoreactive nerve counts normalized to epidermal area, which measures unmyelinated nociceptive nerve fibers in the skin (**Figure 5e**). These findings suggest that SC-specific MCT1 is critical for maintenance of mechanical, but not heat, sensitivity during aging.

### 3.6 Selective ablation of MCT1 impairs glycolytic and mitochondrial functions in SCs

In order to begin to understand how MCT1 deficient SCs have impaired capacity for myelination, their capacity for oxidative metabolism and glycolysis was measured by quantifying the rate of real-time oxygen consumption (OCR; **Figure 6a**) and extracellular acidification (ECAR; **Figure 6b**), respectively, in a live cell assay with the Seahorse extracellular flux analyzer. Given the reduction of the lactate transporter MCT1, which is necessary for transporting lactate produced from glycolysis, it was not surprising that ECAR was reduced. Oligomycin-induced ECAR, an indicator of glycolytic activity, was significantly decreased by MCT1 deficiency in SCs, while ECAR during basal respiration was not changed (**Figure 6c**). Remarkably, both basal oxygen consumption and uncoupled respiration (maximal mitochondrial oxygen consumption capacity following the addition of FCCP, which mimics a physiological “energy demand”) were significantly reduced in cultured SCs with MCT1 deficiency (**Figure 6d**). Spare respiratory capacity (SRC), which is defined as the difference between maximal respiration and basal respiration, is a measure of the ability of the cell to respond to increased energy demand as well as how closely the cell is to respiring at its theoretical maximum. SRC was significantly decreased due to MCT1 deficiency in cultured SCs (**Figure 6e**). These findings demonstrate that MCT1 deficiency impairs glycolytic and mitochondrial functions of SCs and worsens their fitness or flexibility to respond to stress stimuli and/or metabolic demands.

**Figure 6.**
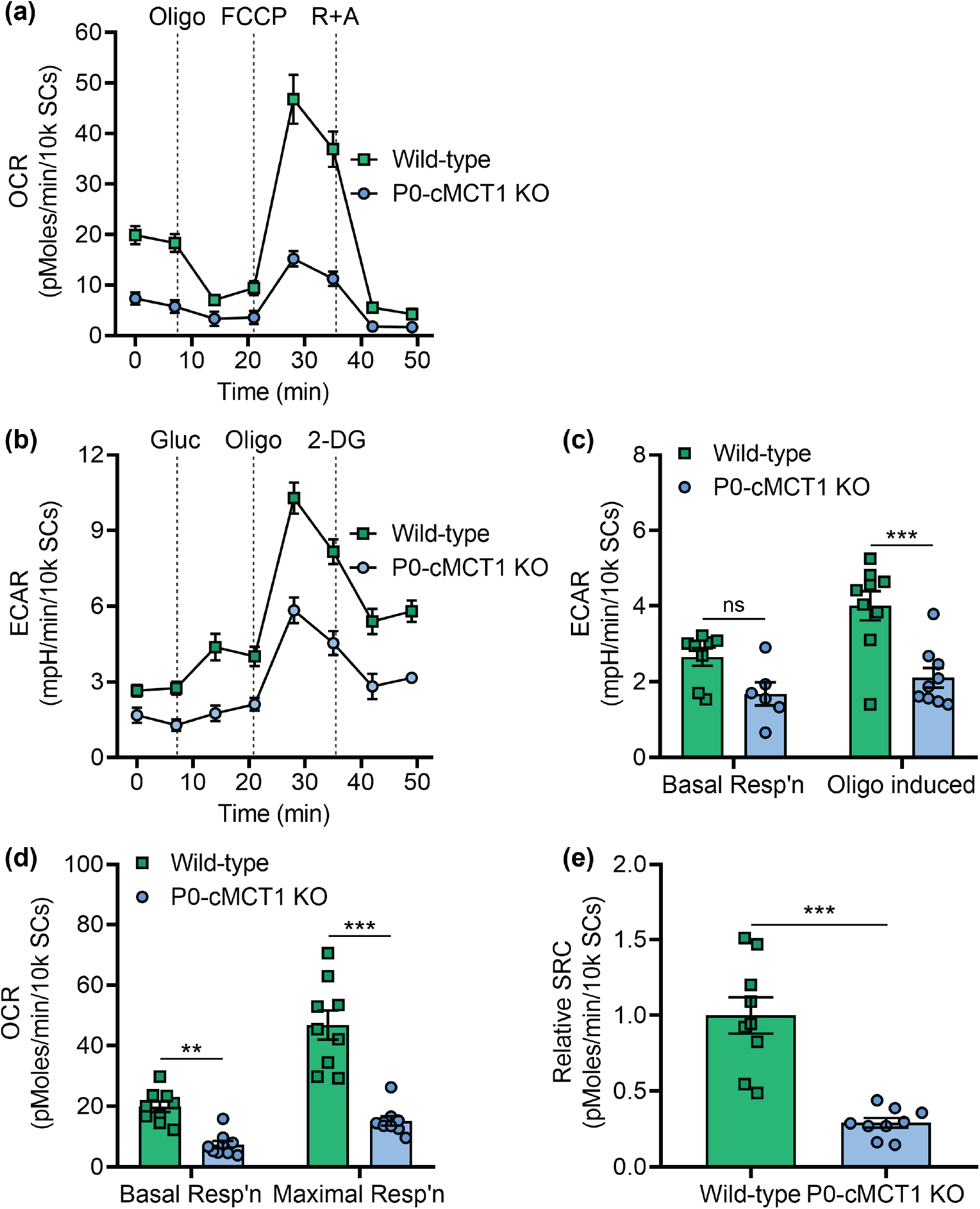
Selective downregulation of MCT1 within SCs impairs glycolytic and mitochondrial functions. Real-time oxygen consumption (**a**) and extracellular acidification (**b**) were measured in SCs isolated from sciatic nerves of 4-week-old wild-type and P0-cMCT1 KO littermate mice with the Seahorse extracellular flux analyzer. The ECAR was compared between wild-type and P0-cMCT1 KO SCs during basal condition and following oligomycin treatment (**c**). The OCR was compared between wild-type and P0-cMCT1 KO SCs during basal respiration and FCCP-induced maximal respiration (**d**). Spare respiratory capacity (SRC) was calculated as the difference between maximal respiration and basal respiration (**e**). Mean ± SEM, *n* = 9 per group, \*\**p* < 0.01; \*\*\**p* < 0.001; ns = not significant, two-way ANOVA with Bonferroni’s multiple comparisons test for C and D; unpaired *t* test for E. ECAR; extracellular acidification rate, OCR; oxygen consumption rate, Oligo; oligomycin, FCCP; carbonyl cyanide-4 (trifluoromethoxy) phenylhydrazone, R+A; Rotenone and Antimycin A, GLUC: glucose, 2-DG: 2-deoxyglucose.

### 3.7 Hypomyelination due to MCT1 deficiency is associated with deficits in synthesis of MAG, but not other myelin proteins, reversion of SC to less mature state, and impaired lipogenesis in peripheral nerves

Several structural proteins play a role in the maintenance of myelin in peripheral nerves. To begin to investigate potential mechanisms for nerve hypomyelination with aging, expression of myelin proteins was investigated in peripheral nerves from aged wild-type and P0-cMCT1 KO mice. Interestingly, expression of major peripheral myelin proteins, myelin protein zero (P0) and myelin basic protein (MBP), were unchanged in sciatic nerves with no MCT1 in SCs (**Figure 7a,b and Supporting Information Figure S7**). In contrast, there was significantly decreased expression of myelin-associated glycoprotein (MAG) from peripheral nerves lacking SC MCT1 (**Figure 7c and Supporting Information Figure S7**). MAG in peripheral nerve is important for long-term stabilization of myelinated axons and prevents axonal degeneration both *in vitro* and *in vivo* (Nguyen et al., 2009). These findings suggest that MCT1 in SCs is critical for synthesis of MAG, but not MBP or P0, during aging.

**Figure 7.**
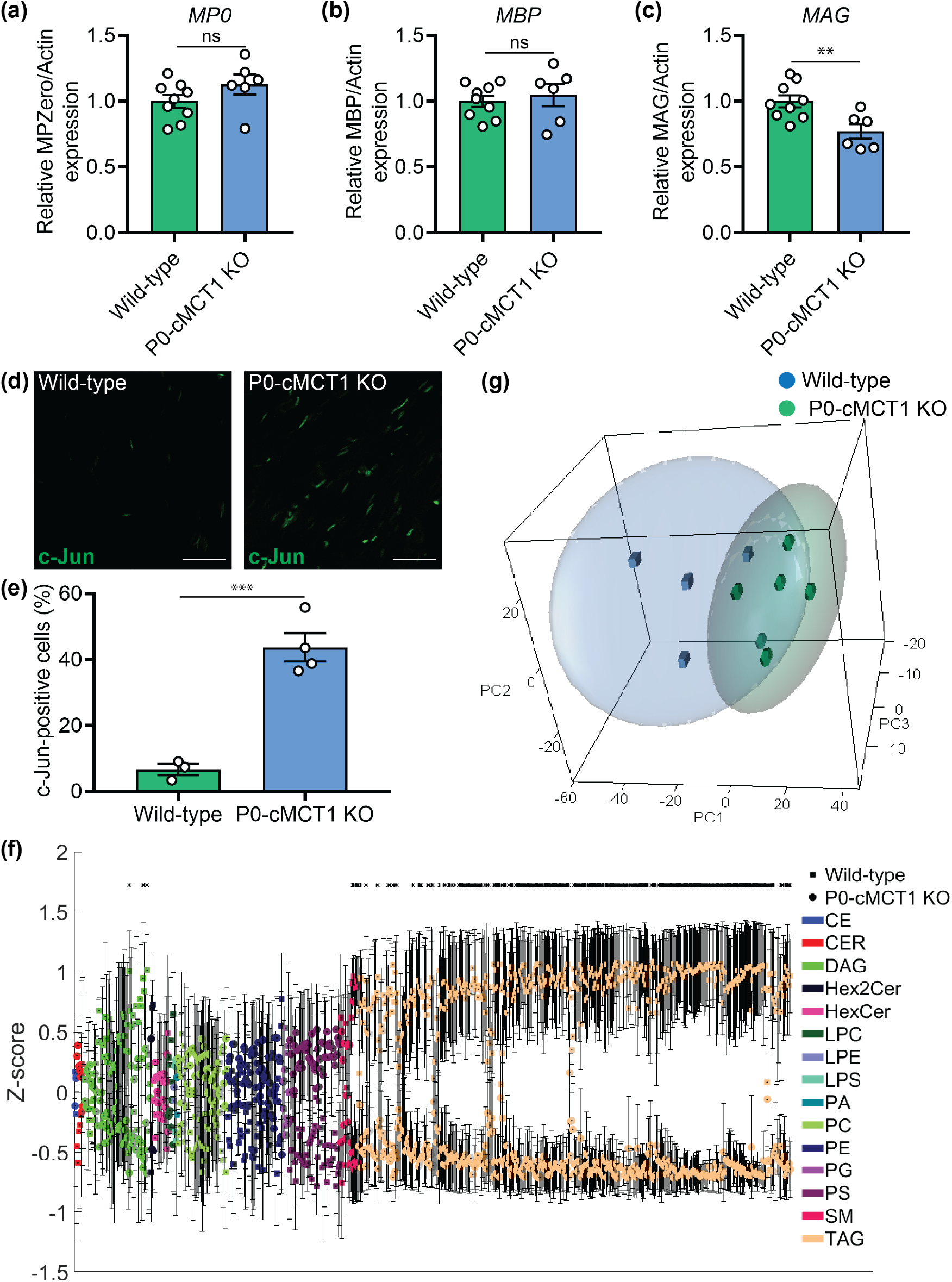
SC-specific MCT1 deficiency decreases the synthesis of myelin-associated glycoprotein (MAG) as well as lipogenesis in peripheral nerves during aging. Expression of myelin protein zero (P0, **a**), myelin basic protein (MBP, **b**), and MAG (**c**) in sciatic nerve homogenates normalized to actin are presented as fold change relative to wild-type control. The full-length Western blots used for the densitometry quantification are shown in **Figure S7**. Mean ± SEM, *n* = 9 for wild-type and 6 for P0-cMCT1 KO groups, \*\**p* < 0.01; ns = not significant, unpaired *t* test. (**d**) Representative images of longitudinal sections of sciatic nerves of mature (four-month-old) wild-type and P0-cMCT1 KO mice immunostained for c-Jun (green). (**e**) Quantification of the c-Jun positive cells in the immunostained longitudinal sections of sciatic nerves presented in panel **d**. c-Jun-positive cells are presented as a percentage relative to the number of DAPI-stained nuclei. Mean ± SEM, *n* = 3 for wild-type and 4 for P0-cMCT1 KO groups, \*\*\**p* < 0.001, unpaired *t* test. (**f, g**) Lipidomics analysis of sural nerves isolated from 12-month old wild-type and P0-cMCT1 KO mice. (**f)** Z-score plot of group means for all detected lipid species. Lipid classes are identified by the indicated color with wild-type = circles, and P0-cMCT1 KO = squares. Please see Supplemental Table 1 for clear identification of individual lipids. Mean ± SEM \**p* < 0.05*, *t* test with Benjamini-Hochberg correction for multiple comparisons. (**g**) Scatter plot of primary component analyses (PCA) scores for wild-type (green spheres) and P0-cMCT1 KO (blue cubes) groups. Ellipses represent 95% confidence intervals. *n* = 4 for wild-type and 6 for P0-cMCT1 KO groups.

The described glycolytic and mitochondrial dysfunction in MCT1-deficient cultured SCs, led us to investigate potential developmental errors in SCs without MCT1. Interestingly, c-Jun expression was dramatically upregulated in sciatic nerves from maturing mutant mice (**Figure 7d,e**). The upregulation of c-Jun, which is a master regulator of SC differentiation (Sasaki, Hackett, Kim, Strickland, & Milbrandt, 2018), in sciatic nerves indicates that MCT1 deficiency causes SCs to arrest at a less mature developmental state. Since the upregulation of c-Jun is also an indicator of mitochondrial stress response (Qureshi, Haynes, & Pellegrino, 2017), the immaturity of SCs due to MCT1 deficiency was further confirmed by upregulated expression of p75-neurotrophin receptor (p75^NTR^), a prototype marker for immature SCs (Chen, Yu, & Strickland, 2007; Jessen & Mirsky, 2008), in sciatic nerves (**Supporting Information Figure S6**).

Myelin is composed of myelin proteins and various lipid species. Given that the two major structural myelin proteins in peripheral nerves, MBP and P0, were unaltered in hypomyelinated nerves from P0-cMCT1 KO mice, the most likely explanation for hypomyelination was reduced synthesis or increased metabolism of SC lipids that are critical components of myelin sheaths. We, therefore, quantified, in an unbiased manner, the abundances of individual lipids from multiple classes in sural nerves from aged mice to explore the influence of SC-specific MCT1 deficiency in lipid metabolism. We detected a total of 717 lipid species by mass spectrometry consisting of 2 cholesterol esters (CE), 8 ceramides (CER), 66 diacylglycerides (DAG), 3 dihexosylceramides (Hex3Cer), 15 hexosylceramides (HexCer), 5 lysophosphatidylcholines (LPC), 2 lysophosphotidylserines (LPS), 1 phosphatidic acid (PA), 50 phosphatidylcholines (PC), 56 phosphatidylethanolamines (PE), 6 phosphatidylglycerols (PG), 51 phosphatidylserines (PS), 18 sphingomyelins (SM), and 433 triacylglycerides (TAG). In univariate analyses, we found that the peak intensities for 382 lipids were significantly different in sural nerves obtained from SC-specific MCT1 deficient mice compared to littermate wild-type mice, and 322 of these lipids remained significant after correction for multiple comparisons (**Figure 7f** and **Supporting Information Table 1**). These lipids consisted of 4 DAGs, 5 SMs, and 313 TAGs. By principal component analysis, the overall PCA explained a cumulative 82.5% of the total variance (with 3 components), and 58.2% of this variance was explained by the first component. There was a visual separation between the lipidomic profiles of sural nerves from wild-type and P0-cMCT1 KO mice in the first three components of the overall PCA (**Figure 7G)**, indicating that loss of myelin from sensory nerves of mutant mice with SC-specific MCT1 deletion is associated with impaired production or increased metabolism of peripheral nerve lipids, especially triacylglycerides, diacylglycerides, and sphingomyelins.

## 4. Discussion

Our laboratory used a newly generated conditional MCT1 knockout mouse to produce mice lacking MCT1in their SCs (i.e., P0-cMCT1 KO mice). To our surprise, MCT1 deletion in SCs led to no compensatory alterations in the expression of other MCTs and GLUTs, and the mice develop normally with no abnormalities in peripheral nerves through two months of age. By four months of age, however, P0-cMCT1 KO mice develop reduced sensory, but not motor, NCV that persists through at least twelve months of age and is associated with a corresponding reduction in myelin thickness and increase in the length of nodes of Ranvier. Myelin thickness and nodes of Ranvier length are the primary structural determinants of the conduction velocity in myelinated axons (Arancibia-Carcamo et al., 2017). This persistent demyelination is associated with impaired sensory function, as measured by mechanical sensitivity, by 12 months of age. These results suggest a functional role of SC MCT1 in sensory nerve myelination and axon biology during aging.

SC metabolism is critical for the regulation and maintenance of peripheral nerve development and function, and SCs are the primary cell-type that metabolically support PNS axons (Beirowski et al., 2014; Jha & Morrison, 2018). Emerging evidence documents a close and regulated metabolic interaction between SCs and neurons in the PNS (Fields, 2015; Jha & Morrison, 2018). GLUTs in endothelial cells (Froehner, Davies, Baldwin, & Lienhard, 1988), perineurial cells (Gerhart & Drewes, 1990; Takebe et al., 2008), and SCs (Magnani et al., 1996) facilitate the uptake of circulating glucose into SCs that can be metabolized to pyruvate, lactate, and other energy intermediates. Our findings show that MCT1 deficiency dramatically alters the bioenergetics of SCs. Given the absence of their primary lactate transporter, it is not surprising that there was a significant decrease in oligomycin-induced ECAR in cultured mutant SCs, indicating impaired glycolytic activity. Interestingly, absence of SC MCT1 limited not only glycolysis, but mitochondrial function, as well. The reason for mitochondrial dysfunction in SCs lacking MCT1 is not clear, but may be due to lactate accumulation and reduced intracellular pH in MCT1-deficient SCs. It should be noted, however, that these metabolic studies were conducted in SC cultures and the toxicity of MCT1 ablation may be accentuated *in vitro* due to specific culture conditions.

Sciatic nerves from maturing P0-MCT1 KO mice have increased expression of both c-Jun and p75^NTR^. c-Jun is a negative regulator of SC differentiation, myelination, and has been implicated in demyelinating neuropathies (Jessen & Mirsky, 2008). p75^NTR^, which has been implicated in the regulation of gene expression, cell cycle transitions, and myelination by recruiting specific binding proteins, is a marker for immature SCs that is highly expressed primarily during development and after nerve injury (Cragnolini & Friedman, 2008; Wang et al., 2017). Though c-Jun is also upregulated by mitochondrial injury, the upregulation of p75^NTR^ confirms the presence of increased immature SCs in P0-MCT1 KO mice. Interestingly, there is a greater increase in c-Jun-positive SCs, as compared to p75^NTR^ –positive SCs, suggesting the presence of mitochondrial injury in SCs *in vivo*. The importance of upregulation of c-Jun and p75^NTR^ is two-fold. First, the upregulation of these proteins suggest that the deficits in myelination are not merely due to “sick” cells with diminished overall metabolism, but rather a shift in the phenotype of these cells to a more premature state. Second, upregulation of c-Jun and demyelinating neuropathy was also recently reported following dysregulation of NAD^+^ biosynthesis, a key cellular factor in glycolytic metabolism (Verdin, 2015). SC-specific loss of MCT1 may similarly mediate c-Jun expression through impairment of glycolysis. Though c-Jun and p75^NTR^ can also be upregulated by nerve injury, this is unlikely the cause in our mice since c-Jun upregulation occurs prior to any evidence of axon injury.

Myelin membranes have a very high lipid-to-protein ratio, in which lipids constitute 70-80% of myelin (O’Brien, Sampson, & Stern, 1967). Lipid biosynthesis is thus critical for SCs during both developmental myelination and maintenance of myelin sheaths during maturation and aging; and abnormalities of lipid metabolism can directly cause hypomyelination (Montani et al., 2018; Schmitt, Castelvetri, & Simons, 2015). Though the two most abundant myelin proteins, P0 and MBP, were present in normal amounts, several lipids, including sphingomyelin, triacylyglycerides, and diacylglycerides were reduced in sensory peripheral nerves lacking SC MCT1. This finding illustrates that SC-specific MCT1 deletion does not significantly affect transcription, translation, or stability of at least the most prominent myelin proteins in peripheral nerves, rather hypomyelination appears to be due to reduced production or increased scavenging of myelin lipids. Sphingomyelin is a major component of peripheral nerve myelin, where it represents 10-35% of the total lipids in mice (Garbay, Heape, Sargueil, & Cassagne, 2000). Degradation of sphingomyelin is reported following peripheral nerve injury (Patti et al., 2012) and chemotherapy-induced peripheral neuropathy (Stockstill et al., 2018); and sphingomyelin content of myelinating DRG cultures correlates with state of demyelination following forskolin treatment (Capodivento et al., 2017). Triacylglycerides are lipids with very short half-lives used primarily for energy production or storage. Though not a component of myelin membranes, themselves, their turnover is quite rapid and they serve as intermediates in the synthesis of longer-chain fatty acids and sphingosine, which are major components of myelin (O’Brien et al., 1967). Interestingly, vitamin B12 deficiency, which causes axonopathy in both spinal cord descending tracts and peripheral nerves (Stabler, 2013), appears to be at least partly dependent on triacylglycerides. Rats on a vitamin B12 deficient diet developed a marked decrease in sciatic nerve triacylglycerides without alterations in other lipids (Turner & Vevallos, 1968). Diacylglyceride is a precursor for triacylglycerides and plays an important role in lipid biosynthesis. In addition, diacylglyceride is itself an important membrane component and key second messenger in multiple cellular signaling cascades (Carrasco & Merida, 2007). Thus, the significant reductions in sphingomyelin, triacylglyceride, and diacylglyceride in P0-cMCT1 KO mice may contribute to the age-dependent demyelination and axon loss in sensory nerves of P0-cMCT1 KO mice. Our hypothesis is that impaired glycolytic and oxidative metabolism in SCs leads to increased utilization and impaired synthesis of SC lipids, thus removing a critical substrate for the maintenance of myelin lipids. Though not previously investigated following abnormalities in MCT1, mitochondrial defects caused reduced pyruvate oxidation, aberrant SC lipid metabolism, and peripheral nerve injury (Riviere et al., 2009); and fatty acid synthesis in SCs is necessary for myelination (Montani et al., 2018). Thus, the observed reduction in mitochondrial oxidative phosphorylation in MCT1-deficient SCs is likely one mechanism for aberrant lipid metabolism. Additionally, reduced glycolysis in MCT1-deficient SCs, by lactate retention reducing glucose uptake and glycolysis and impeding intracellular energy production (Leite et al., 2011), will also reduce the availability of pyruvate for oxidation and fatty acid synthesis, potentially contributing to demyelination.

SC-specific MCT1 ablation causes deficits in sensory, but not in motor, myelination of peripheral nerves by four months of age, as quantified by electrophysiology and histology; and sensory behavioral deficits by 12 months of age. The selective impact on sensory nerves was not due to differential expression of MCT1, evaluated at the level of mRNA, in sensory versus motor SCs. Though the reason for sensory SCs vulnerability remains unknown, emerging evidence shows that SCs associated with sensory or motor nerves have differential functions and distinct molecular phenotypes (Brushart et al., 2013; Hoke et al., 2006; Wright et al., 2014). Our findings add to these phenotypic differences, suggesting that metabolic signatures or susceptibility to metabolic alterations can also differentiate sensory and motor SCs. Interestingly, peripheral neuropathies, particularly those caused by metabolic dysfunction (i.e., diabetes) or toxins, are most commonly sensory predominant (Katona & Weis, 2017). Whether alterations in MCT1 or other metabolic transporters are responsible for this vulnerability has not yet been determined.

Our results indicate that SC MCT1 is critical for maintenance of myelin and, nodes of Ranvier morphology of sensory, but not motor, nerves during aging. Our study details several specific cellular pathways disrupted by deletion of SC MCT1, including MAG expression, SC maturation as reflected by c-Jun and p75^NTR^ expression, and lipid metabolism, which may contribute to hypomyelination and dysfunction of sensory peripheral nerves (**Figure 8**). Further studies manipulating this energetic pathway may reveal potential novel treatments for peripheral neuropathies, many of which are extremely common and currently untreatable.

**Figure 8.**
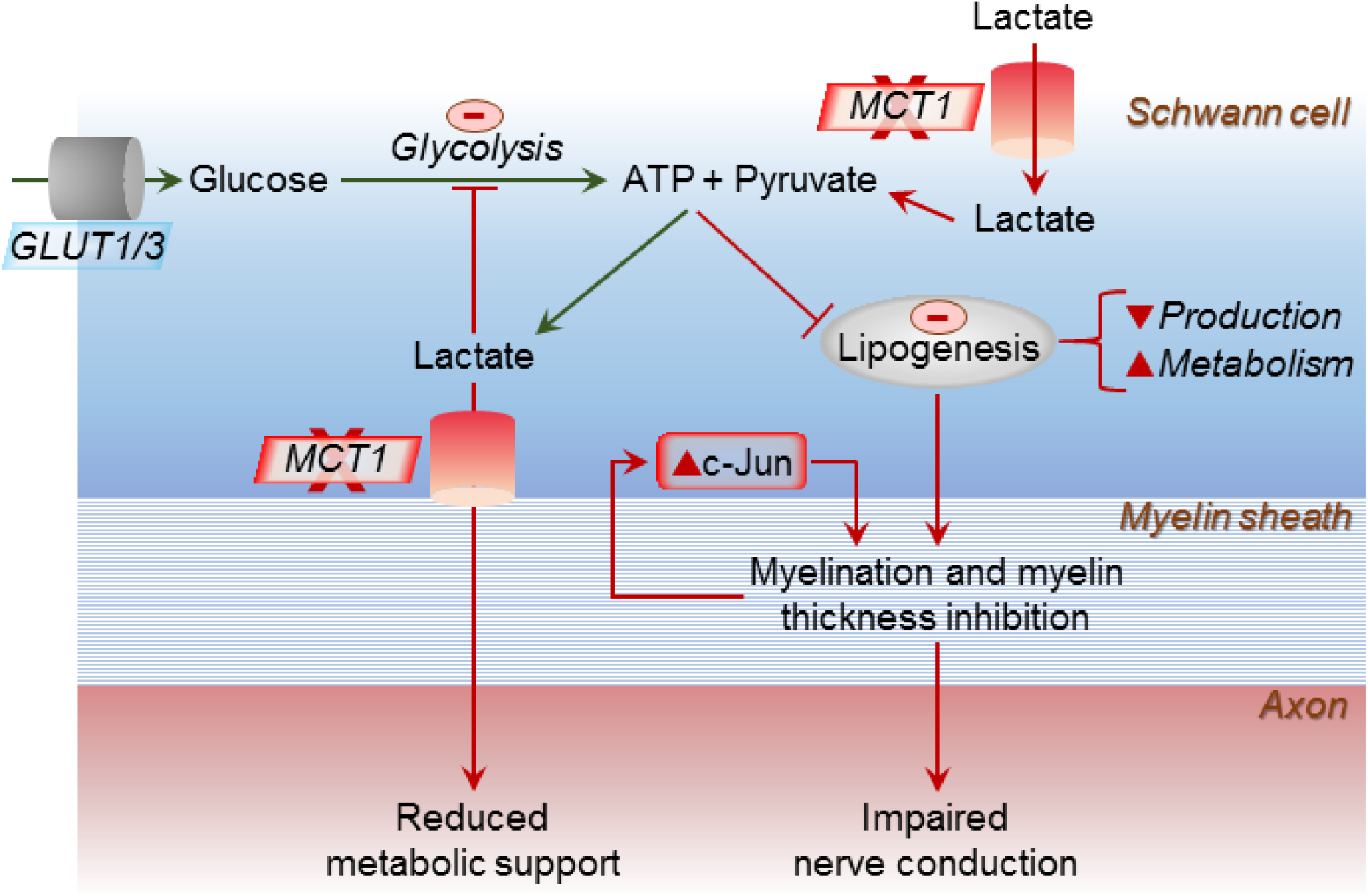
Model for impact of SC ablation of MCT1 on lipogenesis, c-Jun, and axon-myelin function. Circulating glucose is taken up into SCs by glucose transporters, both GLUT1 and GLUT3. Glucose is metabolized to pyruvate and ATP is produced via glycolysis. Besides generating more ATP via oxidative phosphorylation in mitochondria, some of the pyruvate is converted to lactate. Lactate can also be directly imported into SCs from the circulation and extracellular space. This lactate can be exported to axons via MCT1 to generate ATP in their mitochondria or converted to pyruvate for subsequent oxidative phosphorylation in SCs. In the absence of MCT1, both the lactate transport to axons as well as the lactate transport into SCs is disrupted, which can impair lipogenesis, lead to elevated c-Jun, and result in immature, poorly myelinating SCs.

## Acknowledgements

The authors would like to thank Dr. Mohamed Farah, Ms. Carol Cooke, Ms. Kimberly Brown, and the Johns Hopkins Neurology Electron Microscopy Core, for their assistance in processing and photographing electron microscopic pictures and processing embedded nerve tissue for toluidine blue staining. We also thank Dr. Ru-ching Hsia and the University of Maryland Electron Microscopy Core Imaging Facility for assistance with photographing electron microscopic pictures. We would also like to thank the Pain Research Core funded by the Blaustein Fund and the Neurosurgery Pain Research Institute at The Johns Hopkins University for providing facilities for behavioral studies. Financial support was provided by NIH-NS086818-01 (B.M.M.).

## Author Contributions

M.K.J. conceived and performed experiments, analyzed data, and wrote the manuscript. Y.L., K.A.R., F.Y., R.M.D., P.D., X.H.A., W.C., and Y.L. performed some of the experiments and/or data analysis. Y.G., M.J.P., A.H., N.J.H., and J.D.R. assisted with resources. Y.G., A.H., N.J.H., and J.D.R. provided expertise and feedback. B.M.M. secured funding, conceived and supervised the study, analyzed data, and wrote the manuscript.

## Supporting Information

**Figure S1.**
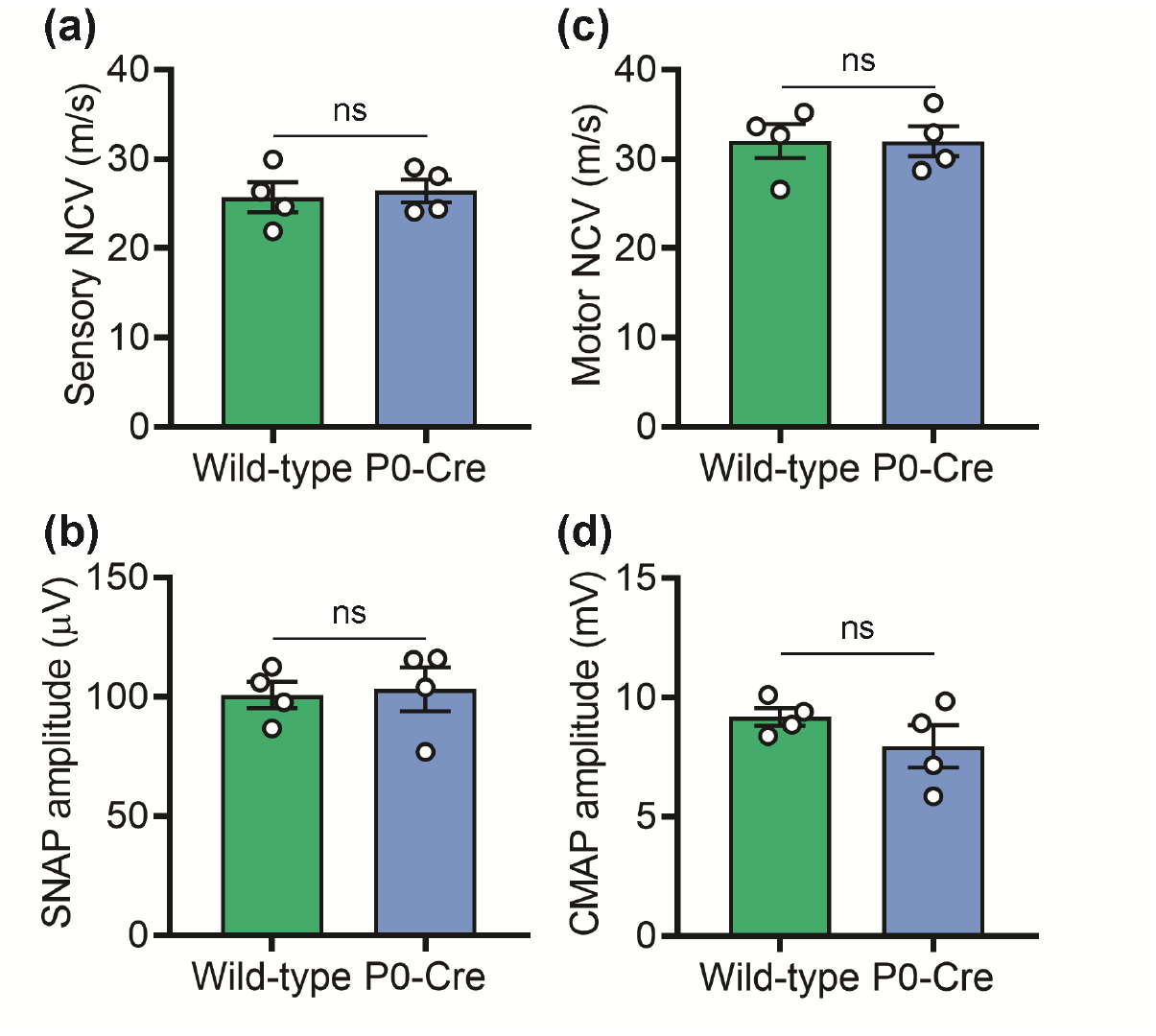
Mice expressing P0-Cre selectively within SCs have unaltered peripheral nerve electrophysiology. (**a-d**) Sensory (**a**) and motor (**c**) nerve conduction velocities are unaltered in mice expressing P0-Cre selectively within SCs (without MCT1*^f/f^*) compared to wild-type littermate control mice. There is also no significant impact on SNAP (**b**) or CMAP (**d**) amplitudes. 4-month-old littermate mice were used for the electrophysiological recordings. Mean ± SEM, *n* = 4 per group, ns = not significant, unpaired *t* test. NCV; nerve conduction velocity, SNAP; sensory nerve action potential, CMAP; compound muscle action potential.

**Figure S2.**
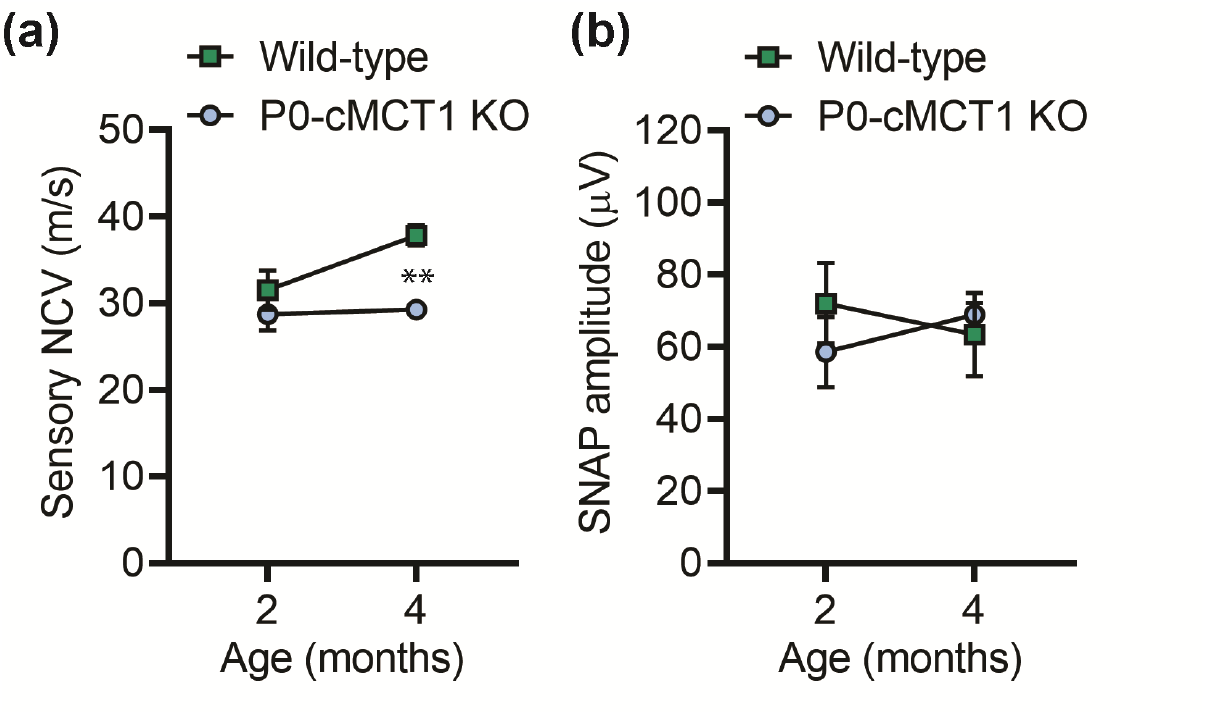
Sensory NCV decreased in maturing P0-cMCT1 KO mice was also evident after warming tail. The decreased sensory NCV (**a**) and unchanged SNAP amplitude (**b**) due to MCT1 deletion in SCs at 4 months of age was also confirmed by the electrophysiologic recordings performed while maintaining tail temperature at 32–34°C. Mean ± SEM, *n* = 4 per group, \*\**p* < 0.01; two-way ANOVA with Holm-Sidak’s multiple comparisons test. NCV; nerve conduction velocity, SNAP; sensory nerve action potential.

**Figure S3.**
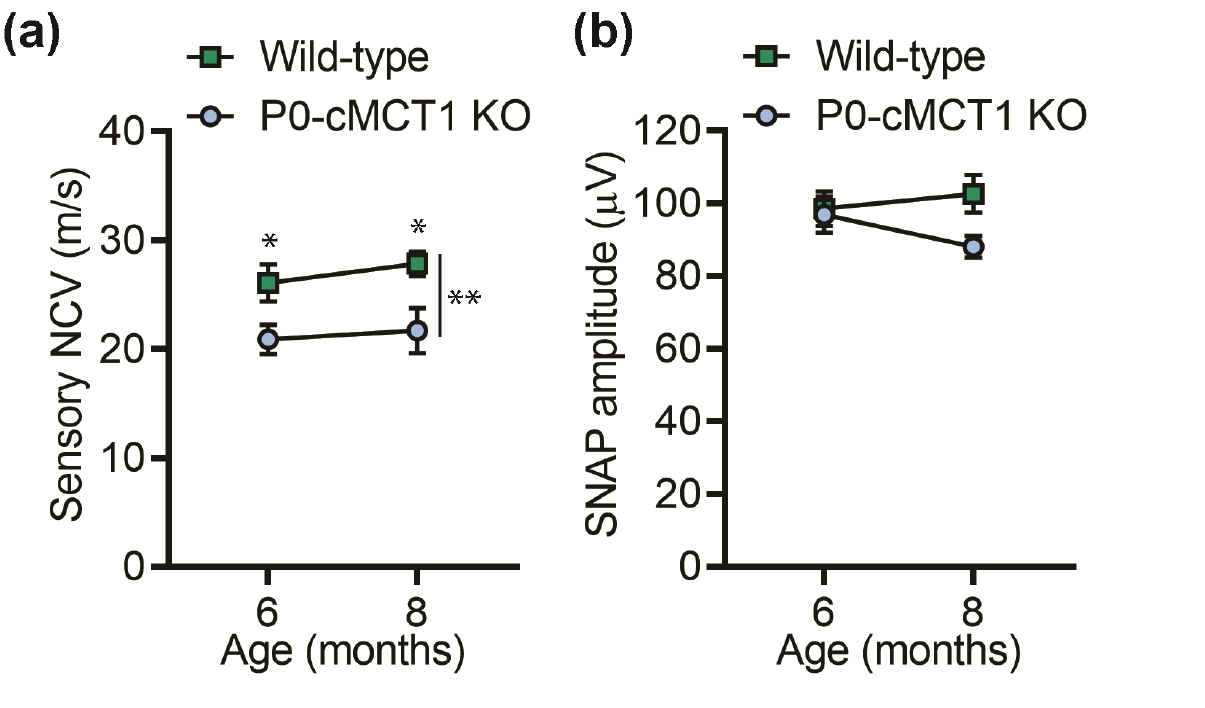
Electrophysiological deficit observed due to MCT1 deficiency in SCs is independent of gender. (**a** and **b**) Assessment of sensory NCV (**a**) and SNAP amplitude (**b**) in male littermate mice having SC-specific MCT1 deficiency. Age-dependent decrease in sensory nerve conduction in female mice due to SC-specific MCT1 deficiency is also evident in male mice. Mean ± SEM, *n* = 5-8 per group per time point, \**p* < 0.05, \*\**p* < 0.01, two-way ANOVA with Holm-Sidak’s multiple comparisons test.

**Figure S4.**
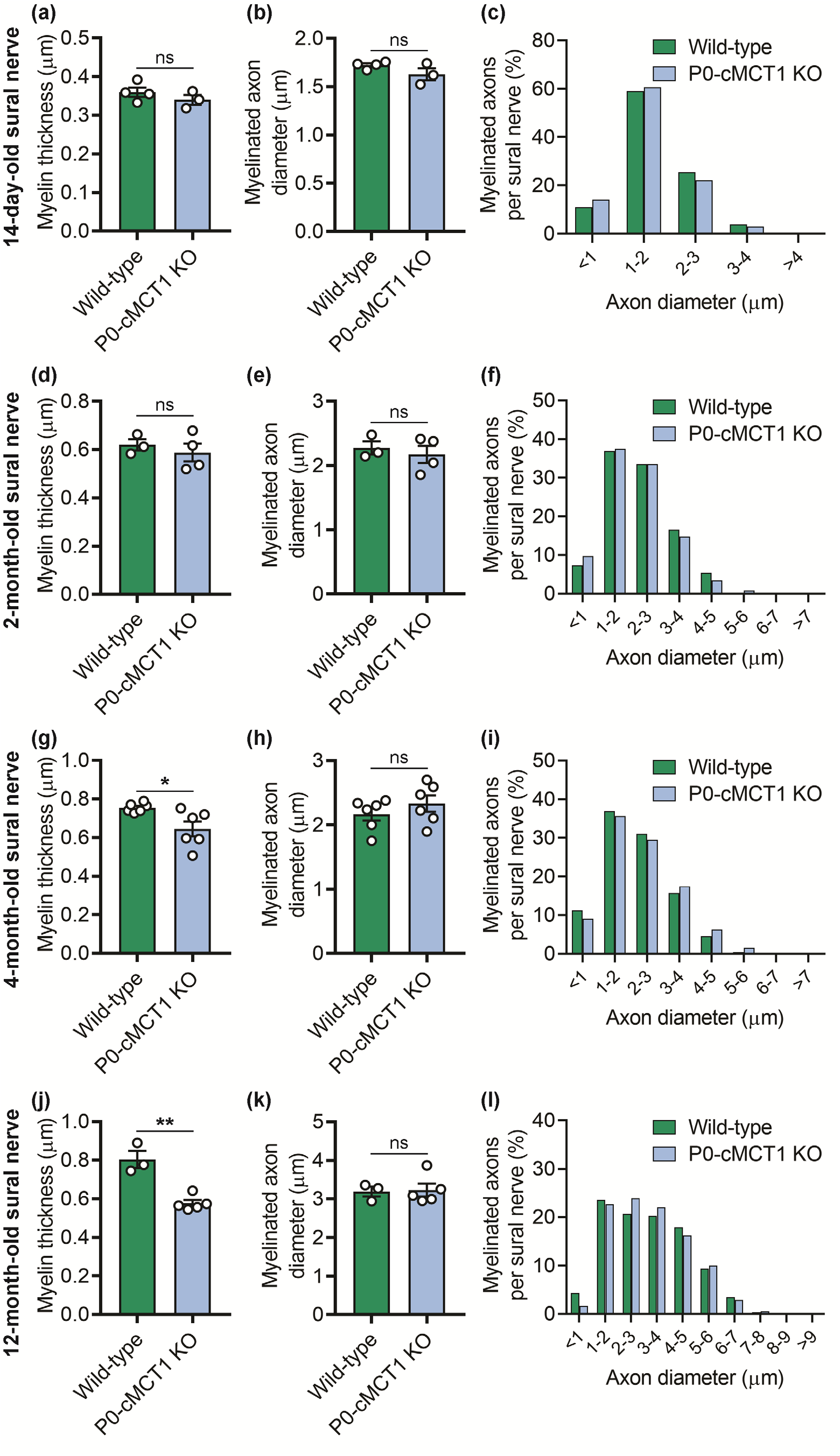
SC-specific MCT1 maintains axon myelin without altering axon caliber in sural nerve during aging. (**a-f**) Morphological analyses of 14-day-old and 2-month-old wild-type and P0-cMCT1 KO mice sural nerves. Myelin thickness (**a,d**), axon diameter (**b,e**), and distribution of myelinated fibers (**c,f**) confirmed normal embryonic and early post-natal sural nerve development in P0-cMCT1 KO mice. (**g-i**) Morphological analyses of 4-month-old wild-type and P0-cMCT1 KO mice sural nerves. Myelin thickness (**g**), axon diameter (**h**), and distribution of myelinated fibers (**i**) confirmed sural nerve hypomyelination without significant alteration in axon caliber in maturing P0-cMCT1 KO mice. (**j-l**) Morphological analyses of 12-month-old wild-type and P0-cMCT1 KO mice sural nerves. Myelin thickness (**j**), axon diameter (**k**), and distribution of myelinated fibers (**l**) confirmed sural nerve hypomyelination without significant alteration in axon caliber in aged P0-cMCT1 KO mice. Light microscope photomicrographs of toluidine blue-stained sections of the sural nerves were analyzed for morphometric analysis. Mean ± SEM, *n* = 3-6 per group, \**p* < 0.05; \*\**p* < 0.01; ns = not significant, unpaired *t* test.

**Figure S5.**
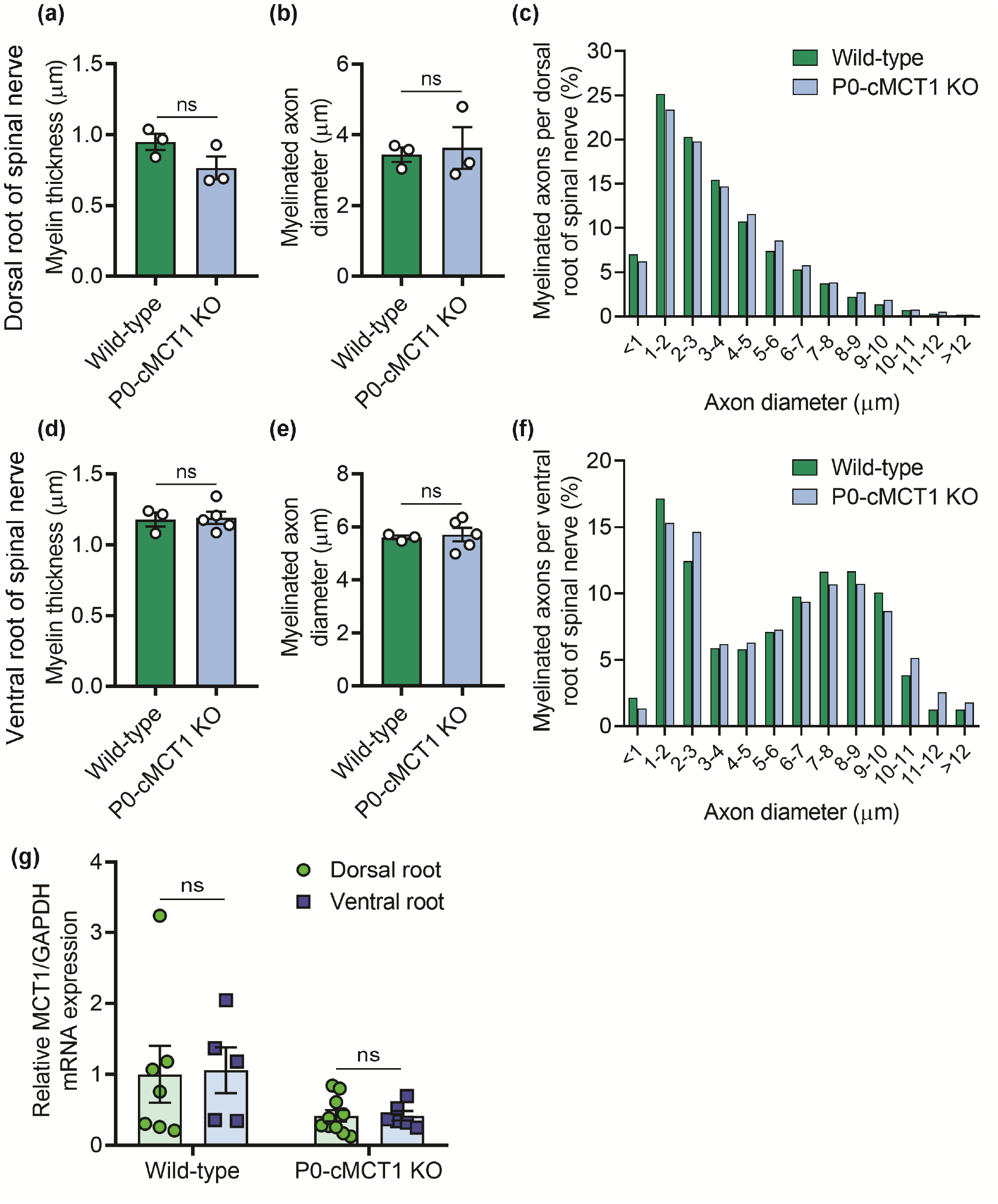
MCT1 expression, overall myelin thickness, and myelinated axon number are similar in both dorsal and ventral roots. (**a-c**) Morphological analyses of 12-month-old wild-type and P0-cMCT1 KO mouse dorsal spinal nerve roots. Myelin thickness (**a**), axon diameter (**b**), and distribution of myelinated fibers (**c**) demonstrated no significant alteration in myelination and axon caliber of dorsal spinal nerve roots in P0-cMCT1 KO mice. (**d-f**) Morphological analyses of 12-month-old wild-type and P0-cMCT1 KO mice ventral spinal nerve roots. Myelin thickness (**d**), axon diameter (**e**), and distribution of myelinated fibers (**f**) demonstrated no remarkable alteration in myelination and axon caliber of ventral root of spinal nerve in P0-cMCT1 KO mice. Light microscope photomicrographs of toluidine blue-stained sections of the dorsal and ventral spinal nerve roots were used for the morphometric analyses. Mean ± SEM, *n* = 3-5 per group, ns = not significant, unpaired *t* test. (**g**) Dorsal and ventral roots of spinal nerves show similar expression of MCT1 mRNA, and SC specific MCT1 deletion downregulates the expression of MCT1 mRNA in both dorsal and ventral roots of spinal nerves. Levels of MCT1 mRNA are depicted as fold change compared with wild-type mice normalized to GAPDH mRNA levels. Mean ± SEM, *n* = 5-10 per group, ns = not significant, two-way ANOVA with Bonferroni’s multiple comparisons test.

**Figure S6.**
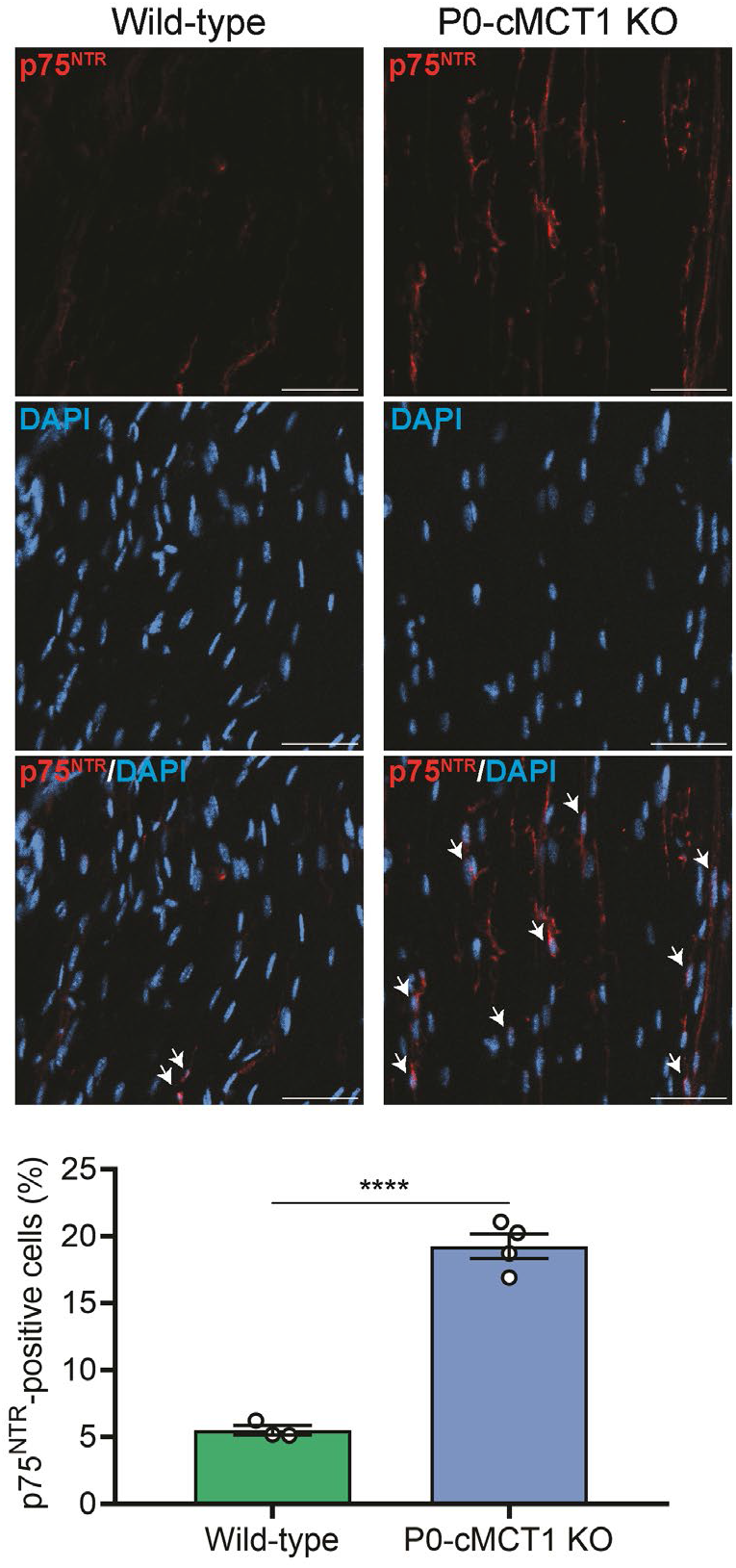
SC-specific MCT1 deficiency increases the expression of p75^NTR^ in peripheral nerves during aging. Representative images (upper panel) of longitudinal sections of sciatic nerves from mature (four-month-old) wild-type and P0-cMCT1 KO mice immunostained with anti-CD271 (p75^NTR^) antibody (red) and DAPI (blue, nuclear counterstain). Examples of co-localization are marked with white arrows in merge panel. Quantification of the p75^NTR^-positive cells in the immunostained longitudinal sections of sciatic nerves is presented in lower panel. p75^NTR^-positive cells are presented as a percentage relative to the number of DAPI-stained nuclei. Mean ± SEM, *n* = 3 for wild-type and 4 for P0-cMCT1 KO groups, \*\*\*\**p* < 0.0001, unpaired *t* test. Scale bars, 50 µm.

**Figure S7.**
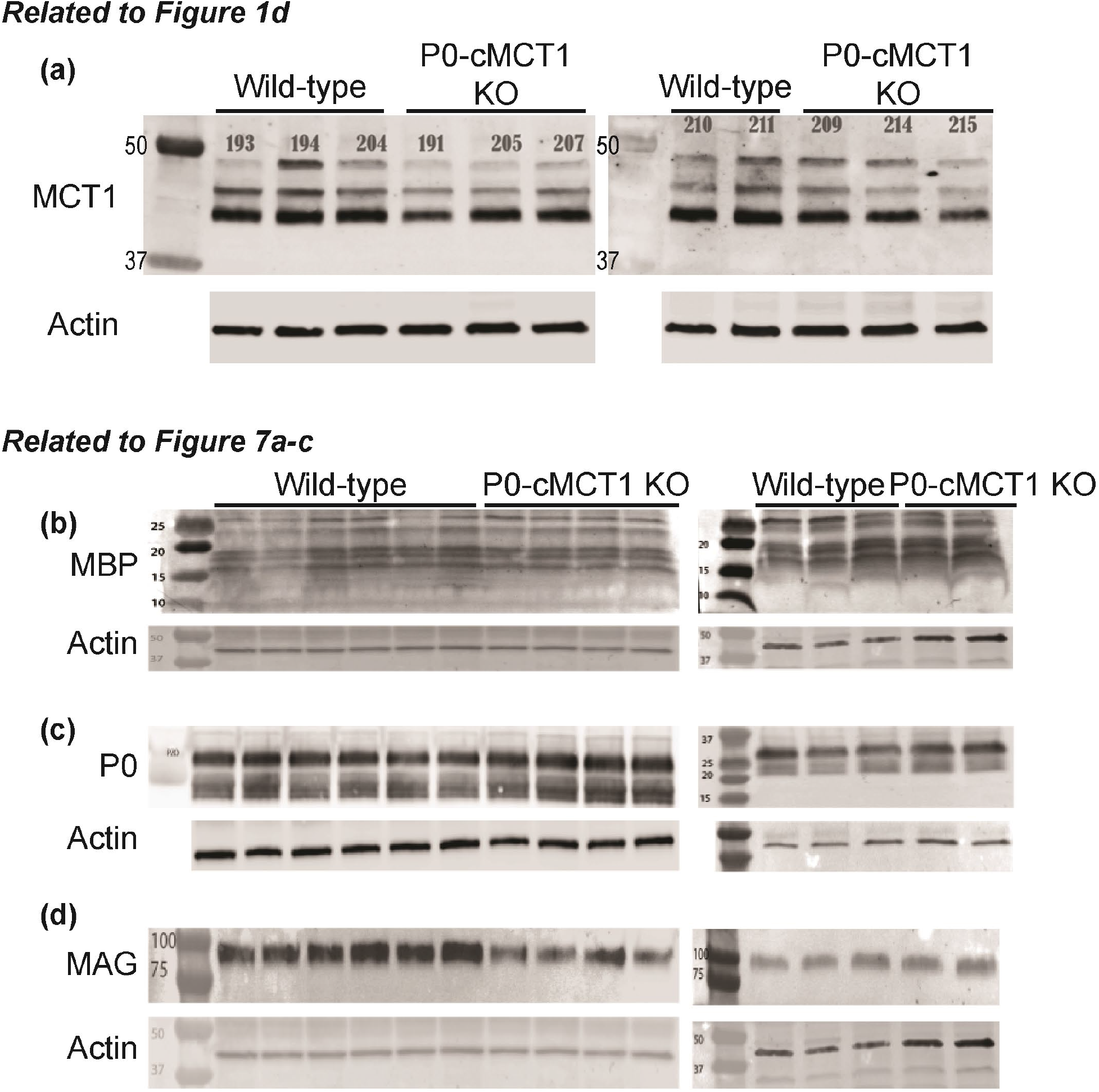
Full-length western blots of MCT1, MBP, P0, and MAG. Full-length Western blots from Figures 1D (**a**) and 7A-D (**b-d**). Protein expression from sciatic nerves of 3 month old (**a**) or 12 month old (**b-d**) wild-type or PO-cMCT1 KO mice for MCT1 (**a**), MBP (**b**), P0 (**c**), and MAG (**d**).

